# Bulk and single-molecule analysis of a novel DNA2-like helicase-nuclease reveals a single-stranded DNA looping motor

**DOI:** 10.1101/851329

**Authors:** O.J. Wilkinson, C. Carrasco, C. Aicart-Ramos, F. Moreno-Herrero, M.S. Dillingham

## Abstract

DNA2 is an essential enzyme involved in DNA replication and repair in eukaryotes. In a search for homologues of this protein, we identified and characterised *Geobacillus stearothermophilus* Bad, a novel bacterial DNA helicase-nuclease with similarity to human DNA2. We show that Bad contains an Fe-S cluster and identify four cysteine residues that are likely to co-ordinate the cluster by analogy to DNA2. The purified enzyme specifically recognises ss-dsDNA junctions and possesses ssDNA-dependent ATPase, ssDNA binding, ssDNA endonuclease, 5’ to 3’ ssDNA translocase and 5’ to 3’ helicase activity. Single molecule analysis reveals that Bad is a highly processive DNA motor capable of moving along DNA for distances of more than 4 kbp at a rate of ∼200 base pairs per second at room temperature. Interestingly, as reported for the homologous human and yeast DNA2 proteins, the DNA unwinding activity of Bad is cryptic and can be unmasked by inactivating the intrinsic nuclease activity. Strikingly, our experiments also show that the enzyme loops DNA while translocating, which is an emerging feature of highly processive DNA unwinding enzymes. The bacterial Bad enzymes will provide an excellent model system for understanding the biochemical properties of DNA2-like helicase-nucleases and DNA looping motor proteins in general.

## INTRODUCTION

DNA2 is an essential replication and repair factor found widely in eukaryotic genomes (1). It is a multi-functional protein with roles in the processing of Okazaki fragments, stalled replication forks, and double-stranded DNA break repair (2–6).The primary structure of DNA2 comprises an N-terminal RecB-family nuclease domain fused to a C-terminal SF1B helicase domain (7). In previous work, we hypothesised that the nuclease domain found in DNA2 and closely-related enzymes belonged to a new class of 4Fe-4S cluster associated domains (8), and this was later confirmed experimentally (9). We called these “iron-staple” nuclease domains because of a unique arrangement of the cysteine residues that co-ordinate the cluster (8, 10). Interestingly, we found that iron-staple domains were very commonly found associated with Superfamily I helicase domains, and further examples included components of some CRISPR systems (11). Given that DNA2 was thought to be restricted to eukaryotic and archaeal organisms, we were surprised to also find DNA2-like enzymes sporadically distributed in restricted niches of bacteria including *Geobacilli* and *Mycobacteria* (see **Supplementary Figure 1** for further information).

From a mechanistic viewpoint, DNA2-like enzymes may be regarded as somewhat peculiar. Structural and biochemical analyses have shown that this domain arrangement places an endonuclease activity “in front” of a translocating motor, such that the enzyme appears to have evolved to cleave the DNA track ahead of itself as it moves along DNA (12, 13). In accordance with this proposition, the nuclease activity has been shown to be inhibitory to the translocase activity *in vitro* (12, 14). This contrasts with the more intuitive and common domain arrangement seen in many other helicase-nuclease fusions, where the nuclease domain is transported behind the translocating motor, potentially leading to processive DNA degradation activity (15–17). To learn more about the activity of this intriguing class of proteins we have cloned, expressed and purified Bad, a <u>ba</u>cterial <u>D</u>NA2-like enzyme from *Geobacillus stearothermophilus*. We show here that Bad contains an Fe-S cluster and identify the four cysteine residues that are likely to co-ordinate the co-factor. The purified enzyme possesses ssDNA binding, ssDNA-dependent ATPase, ssDNA endonuclease, 5’-to-3’ ssDNA translocase and 5’-to-3’ helicase activity. Single molecule analysis reveals that Bad acts as a fast and highly processive DNA looping motor, but that this activity is only evident under conditions in which the nuclease activity is suppressed, either by mutagenesis or by reducing the concentration of free Mg^2+^ ions in the reaction buffer. Thus, Bad displays highly similar biochemical properties to its homologues (including the human and yeast DNA2 proteins) and provides a robust model system for the study of DNA-looping helicases using single-molecule analysis.

## MATERIALS AND METHODS

### Identification, cloning and expression of Bad, a <u>ba</u>cterial <u>D</u>NA2-like enzyme

Novel DNA2-like enzymes were identified by searching for uncharacterised proteins containing motifs characteristic of Superfamily I DNA helicases and Fe-S-containing nuclease domains (see **Supplementary Figure 1-3** for a comparison of Bad with the canonical DNA2 enzymes from yeast and human cells). A DNA2-like enzyme from *Geobacillus stearothermophilus 10* (DSM accession number 13240, Taxonomy ID: 272567) was cloned by PCR from genomic DNA using standard techniques. The untagged gene was ligated into the pET28a vector (Novagen) for expression using the T7 promoter system. The entire sequence of the cloned gene, which has been annotated as a AAA+ ATPase, was identical to that reported in the *G. stearothermophilus 10* genome (accession number WP_053413574). The gene encodes a 1245 amino acid protein with a molecular weight of 143 kDa and a theoretical extinction co-efficient of 195960 M^-1^ cm^-1^. We note that a small number of other polypeptide sequences have been reported in the proteomes of *Geobacillus stearothermophilus* and closely-related strains which are virtually identical to Bad but which feature an additional 25 amino acids at the N-terminus (e.g. accession number ADU93794.1). Given that sequences of this type are in a significant minority, and that the longer sequences all contain a methionine residue at position 26, it is possible that the start codon has been mis-assigned in these instances. We have also cloned and expressed an example of the putative longer Bad polypeptide, but this was found to be insoluble upon expression in *E. coli* (data not shown).

### Protein Purification

Wild type and mutant Bad proteins were all overexpressed in *E. coli* using the same method. pET28a-Bad was transformed into BL21(DE3) cells. Cells were grown in LB to mid-log phase before induction with IPTG (1 mM) for 16h at 27°C. All buffers were degassed extensively prior to use in order to help prevent oxidation of the putative iron-sulphur cluster. The pellet (∼10g) from 4L of bacterial culture was resuspended in 30 mL lysis buffer (50 mM Tris-HCl pH 8.3, 1 mM EDTA, 150 mM NaCl, 10% glycerol, 5 mM DTT, 1 mM PMSF, and Roche protease inhibitor cocktail) and sonicated on ice. After centrifugation at ∼50,000g for 30 min at 4°C, ammonium sulphate was added slowly with stirring to the cleared lysate to a final concentration of 50% (w/v) at 4°C. The precipitated protein was then recovered to a pellet by centrifugation at 50,000g for 30 min at 4°C. This pellet was then resuspended in Buffer A (20 mM Tris-HCl pH8.0, 1 mM EDTA, 5% glycerol, 0.1 mM PMSF, 5 mM DTT, Roche protease inhibitor cocktail) up to a volume where the conductivity of the solution was 16 mSv, and then loaded at 2 mL/min onto a 5 mL Heparin column equilibrated in Buffer B (20 mM Tris-HCl pH 8.0, 1 mM EDTA, 100 mM NaCl, 5% glycerol, 0.1 mM PMSF, 5 mM DTT). After washing for 5 column volumes with Buffer B, a gradient was run over 20 column volumes into Buffer C (20 mM Tris-HCl pH 8.0, 1 mM EDTA, 1 M NaCl, 5% glycerol, 0.1 mM PMSF, 5 mM DTT). The Bad-containing fractions were pooled and diluted with Buffer A to give a final salt concentration of ∼170 mM NaCl before loading at 2mL/min onto a 1 mL MonoQ column pre-equilibrated in Buffer D (20 mM Tris-HCl pH 8.0, 100 mM NaCl, 5 mM DTT). After washing with 5 column volumes Buffer D, the protein was eluted by running a gradient from Buffer D to 50% Buffer E (20mM Tris-HCl pH 8.0, 1 M NaCl, 5 mM DTT) over 20CV. The most concentrated Bad-containing fractions were pooled and 1.5 mL was injected onto a pre-equilibrated Superdex 200 16/600 column in Buffer F (20 mM Tris-HCl pH 8.0, 300 mM NaCl, 5 mM DTT). The pool from the peak fractions was spin concentrated (Millipore, 50kDa cut-off) to a final concentration of ∼15 µM. *E. coli* SSB protein was expressed and purified as described previously (18) which is a modification of the method developed by Lohman (19).

### Iron chelation assay

The iron content of Bad preparations was determined using bathophenantroline, which chelates ferrous iron (Fe^2+^) resulting in the appearance of an absorbance peak at 535nm, as described previously (8). Briefly, 10 µL of a 15 µM solution of Bad was mixed with 3 µL conc. HCl and incubated at 100°C for 15 min to denature the protein. After centrifugation at 13,000g for 15 min, the supernatant was removed and neutralised with 130 µL of 0.5M Tris-HCl pH 8.5 and then treated with ascorbic acid to a final concentration of 0.26% to reduce the iron. After addition of bathophenanthroline disulphonic acid disodium salt to 0.021%, the samples were incubated for 1h at room temperature and then the absorbance measured at 535nm. The concentration of iron was calculated using the extinction coefficient 22,369 M^-1^cm^-1^.

### ATPase assay

ATPase activity was measured by coupling the hydrolysis of ATP to the oxidation of NADH which gives a change in absorbance at 340 nm. Reactions were performed in a buffer containing 20 mM Tris-Cl pH 8.0, 50 mM NaCl, 5 mM DTT, 1 mM MgCl_2_, 50 U/mL lactate dehydrogenase, 50 U/mL pyruvate dehydrogenase, 1 mM PEP and 100 µg/mL NADH. Rates of ATP hydrolysis were measured over 1 min at 25°C. For calculation of K_DNA_ (defined as the concentration of DNA at which ATP hydrolysis is half-maximal), the ATP concentration was fixed at 2 mM. The Michaelis-Menten plot was performed at 20 μM DNA (nucleotide concentration) which is approximately 10x the K_DNA_ value. The concentration of Bad was 10 nM in these assays unless indicated otherwise.

### Nuclease assay

80 µM (nucleotides) ϕX174 Virion DNA in 50 mM Tris-HCl pH7.5, 30 mM NaCl, 2 mM MgCl_2_, 2 mM DTT, 0.2 mg/mL BSA was treated with 1µM Bad. 10µL aliquots were quenched at time intervals over 60 min with an equal volume of stop buffer (100 mM EDTA, 1% SDS, 15% glycerol) and loaded onto a 1% agarose 1xTAE gel. The gels were stained with ethidium bromide and visualised by UV.

### Streptavidin displacement assay

Streptavidin displacement assays were based on the method of Morris and Raney (20), modified as in (21). 5nM (molecules) of 5′-^32^P-labelled substrate oligonucleotides were incubated with 400 nM streptavidin in 25 mM HEPES pH 7.8, 25 mM NaCl, 2 mM MgCl_2_. Substrates were modified with either a 5′ or 3′ biotin moiety as indicated. The reaction was initiated by adding an equal volume of protein solution in the same buffer to give final concentrations of 10 nM Bad, 5 mM ATP and 8 µM biotin. The reaction was incubated at 37°C and stopped at certain points within a 4 min time course by quenching with an equal volume of stop buffer (300 mM EDTA, 400 mM NaCl, 30 µM poly(dT)). The products were separated on 10% polyacrylamide 1xTBE gels and visualised by phosphorimaging using a Typhoon imager. The sequences of the oligonucleotides used in this assay can be found in the **Supplementary Methods**.

### Helicase assay

Strand-displacement assays were based on the method of Matson (22), modified as in (21). 1 nM (molecules) of 5′-^32^P-labelled substrate oligonucleotides were incubated with 5 nM Bad in either 20 mM Tris-HCl pH7.5, 2 mM MgCl_2_, 3 mM ATP, 1 mM DTT (which we refer to as the “low magnesium” condition) or 20 mM Tris-HCl pH7.5, 4 mM MgCl_2_, 2 mM ATP, 1 mM DTT (which we refer to as the “high magnesium” condition) for 5 min at 25°C. The reaction was quenched at certain intervals over the time course by adding an equal volume of stop buffer (200 mM EDTA, 1% SDS, 10% (w/v) Ficoll 400 and 100 nM of an unlabelled form of the radiolabelled strand in the substrate to prevent re-annealing. The products were separated on 15% polyacrylamide 1xTBE gels and visualised by phosphorimaging using a Typhoon imager. The sequences of the oligonucleotides used in this assay can be found in the **Supplementary Methods**.

### Magnetic Tweezers translocation assay

We used a Magnetic Tweezers setup similar to one reported previously (23). Raw data was recorded at 60 Hz and filtered to 3 Hz for representation and analysis. Force values were calculated using the Brownian motion method applied to a DNA-tethered bead (24). The fluidic chamber was pre-incubated with 0.1 mg ml^-1^ of BSA proteins to minimize non-specific attachments of proteins and beads with the surface. DNA substrates (**Figures 6 and 11**) essentially consist of a DNA molecule of ∼6.6 kbp containing a flap sequence (poly-dT oligo of 37 nt) in a specific-site, and flanked by two smaller fragments (∼1 or 0.6 kb) that act as the immobilisation handles as they are labeled with biotins or digoxigenins. The labeled parts are used to specifically bind each DNA end to a glass surface covered by anti-digoxigenins and to streptavidin coated magnetic beads. MT2 also contains a nick in a specific position within the top (DNA-nick top) or bottom (DNA-nick bottom) strand (**Figure 11**). Doubly-tethered beads were identified by applying magnet rotation on the beads and not considered for the analysis. Unless indicated otherwise, single-molecule translocation experiments were carried out at room temperature and at 8 pN or 14 pN as indicated, in a buffer that contained 20 mM Tris-HCl pH 8.0, 30 mM NaCl, 2 mM MgCl_2_, 5 mM DTT, 4 mM ATP and 100 µg ml^-1^ BSA (i.e. “low magnesium” conditions) with Bad proteins at the quoted concentrations (30 nM, 50 nM, or 163 nM). To initiate the reaction, Bad was flowed into the fluid chamber at 20 μl/min while the positions of the beads were measured in real-time by video microscopy. A fluidic chamber made with one parafilm layer (50 µl total volume) and vertical alignment magnets with a 0.11 mm gap were used to reach high applied forces. The quoted distances in base pairs were corrected using the value given by the worm-like chain model of rise per base pair of dsDNA at a given force. The unwinding rate was calculated by using the derivative of the smoothed data at 3 Hz in order to separate movement from pausing events (23).

## RESULTS

### Identification and purification of a bacterial DNA2-like enzyme

DNA2 is a DNA helicase-nuclease that is ubiquitous in eukaryotic cells and has been shown to be an essential DNA replication and repair factor (1). It is characterised by the fusion of a specific subtype of the RecB-family nuclease domain that contains a 4Fe-4S cluster (the “iron staple” nuclease domain(8)) to a C-terminal SF1B helicase domain. In previous work, we identified and characterised the first example of an iron staple nuclease domain in the AddB subunit of the bacterial enzyme AddAB which, like DNA2, is implicated in the resection of double-stranded DNA breaks (8, 25). Iron staple nuclease domains seem to be rare in nature but they are easily identifiable using a bioinformatics approach. In addition to four amino acid motifs associated with nuclease activity that are shared by all members of the RecB nuclease family, they also contain four strictly conserved Cys residues in a unique pattern that spans the entire domain (8, 10). This arrangement results in the Fe-S cluster being critical for the overall structural integrity of the domain, at least in the case of AddAB. We used this bioinformatics signature to predict Fe-S nuclease domains in other proteins. Prominently, these included DNA2 and the Cas4 enzyme from CRISPR-Cas systems (9, 11). Since DNA2 had been considered a eukaryotic protein, we were surprised to find that our searches also uncovered a few examples of bacterial and archaeal enzymes that displayed a DNA2-like domain architecture (**Supplementary Figures 1-3**). Although these <u>ba</u>cterial <u>D</u>NA2-like (Bad) proteins number relatively few and are found sporadically in the bacterial family tree, they are nevertheless broadly distributed. For example, DNA2-like proteins that are clearly homologous are found both in the Firmicute division of Gram-positive organisms and in some Gram-negative Proteobacteria (see **Supplementary Figure 1**). A second class of bacterial DNA2-like protein is also found in *Mycobacteria* and related organisms including *Rhodococcus*. Finally, similar proteins are also found in Euryarchaea including *Methanobacteria* (data not shown).

To investigate the properties of these enigmatic enzymes we cloned and purified the DNA2-like enzyme Bad from *Geobacillus stearothermophilus*. The protein was well expressed in *E. coli* and purified to near homogeneity without the use of tags (**Figure 1A**). SEC-MALS analysis of the wild type protein in the absence of ligands showed that Bad is a monomeric protein (**Figure 1B**). Purified Bad displayed a golden yellow colour, characteristic of Fe-S cluster containing proteins, and bathophenanthroline assays showed the presence of approximately 3 moles of iron per mole of protein in the preparation (**Figures 1C and 1D**). Given the propensity of Fe-S clusters to be lost during purification, and the primary structure similarity with the AddB nuclease domain (for which the Fe-S is well characterised), it is highly likely that the protein in fact contains a 4Fe-4S cluster (25). However, based on these data we cannot exclude the idea that the protein contains a different type of cluster. Mutation of the cysteine residues that are predicted to co-ordinate the Fe-S cluster (to alanine) resulted in the production of labile protein that was lost during purification (data not shown), presumably because of unfavourable effects on folding. In addition to these Cys to Ala mutations, we also altered amino acids in helicase motif I (i.e. the Walker A motif; K815A) and nuclease motif III (D150A) to generate proteins that would be expected to be devoid of ATPase and nuclease activity, respectively, for use in later experiments (26, 27). The resulting mutants were well-expressed and purified to homogeneity in the same manner as the wild type (**data not shown**).

**Figure 1.**
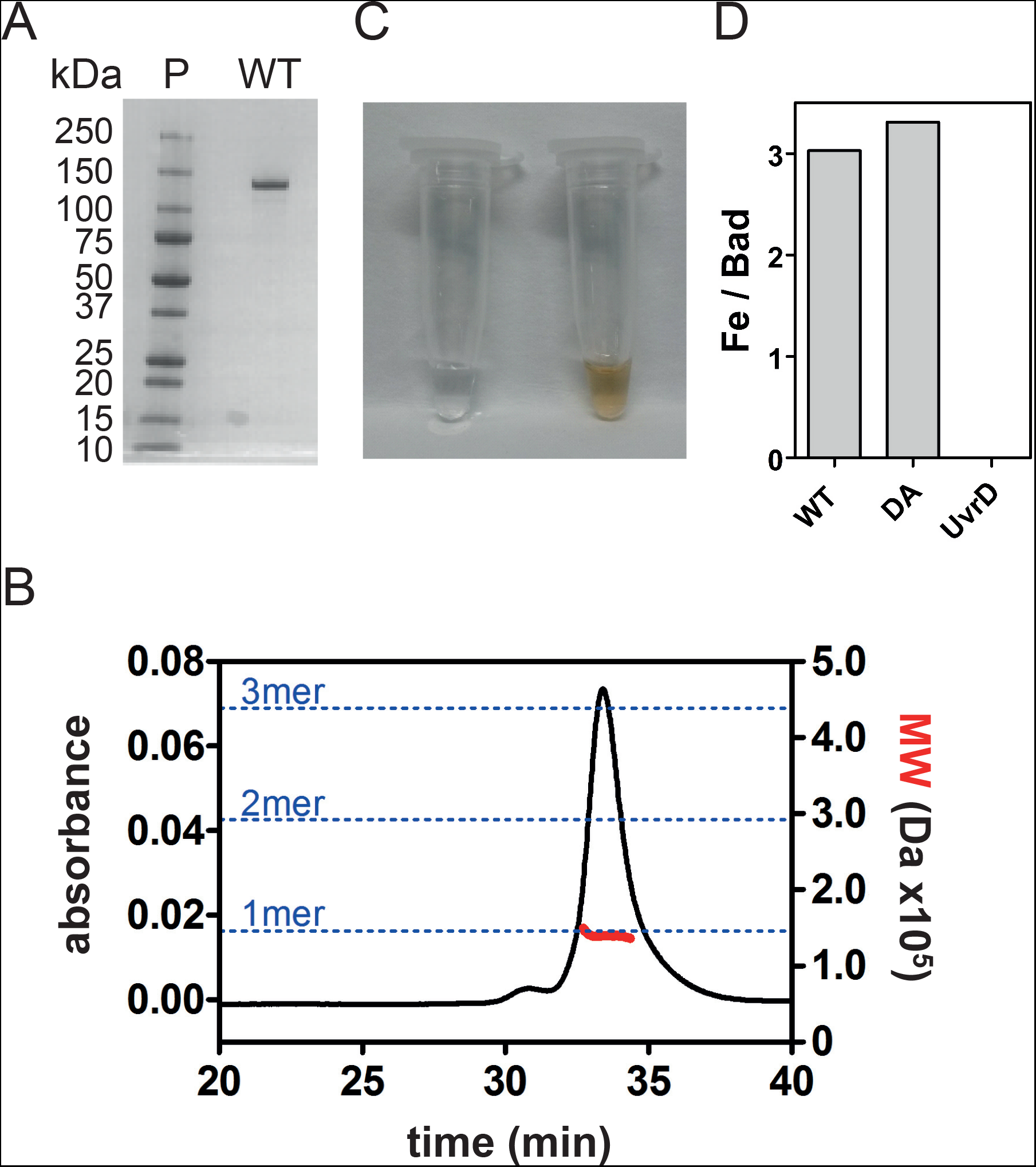
Purification of Bad; a DNA2 homologue from *Geobacillus stearothermophilus*. (A) SDS-PAGE gel showing wild type and mutant Bad proteins used in this study. (B) SEC-MALS analysis of wild type Bad suggests that Bad is a monomer in free solution. (C) The concentrated wild type Bad protein has a golden yellow colour which is characteristic of Fe-S proteins. Storage buffer is shown for comparison. (D) Bathophenanthroline assay reveals the presence of approximately 3 moles of iron per mole of protein in Bad preparations. Data for wild type (WT) and nuclease-dead mutant (DA) Bad are shown, alongside a negative control experiment using *E. coli* UvrD helicase which does not possess an Fe-S cluster.

### Bad is a ssDNA-dependent ATPase and ATP-independent endonuclease

In the presence of saturating quantities of ssDNA, Bad hydrolysed ATP with Michaelis-Menten kinetics, yielding k_cat_ = 101 s^-1^ and K_m_ (ATP) = 220 µM (**Figure 2A**). The turnover number was unaffected by Bad concentration over a wide range of concentrations, providing no evidence for protein association affecting ATPase activity, and consistent with the idea that the protein is functional as a monomer (**Figure 2B**). In the absence of DNA, purified Bad displayed a basal ATPase turnover rate of ∼1 s^-1^ (**Figures 2C and 2D**). In the presence of saturating ATP, titrations with single-stranded DNA revealed half maximal stimulation at K_DNA_ = 1.6 µM nucleotides (**Figure 2C**). Activation of the ATPase activity was apparent regardless of whether the ssDNA was linear or circular, whereas dsDNA was a relatively poor cofactor (**Figure 2D**). These properties are typical of the Superfamily I helicases of which Bad is a member (27). Mutation of helicase motif I (K815A, also known as the Walker A motif) dramatically decreased the ATPase activity showing that it is intrinsic to the Bad polypeptide, whereas mutation of the nuclease motif (D150A) had little effect on the steady-state ATPase activity (**Figure 2E**). We next investigated the nuclease activity associated with Bad in the absence of ATP. To test this, we incubated Bad with circular ssDNA in the presence or absence of Mg^2+^ ions which would be expected to be required for activity. Bad was able to endonucleolytically cleave the ssDNA in a Mg^2+^-dependent fashion (**Figure 2F**). Mutation of nuclease motif III (Bad D150A) eliminated the observed DNA degradation, demonstrating that this activity is also intrinsic to the Bad polypeptide.

**Figure 2.**
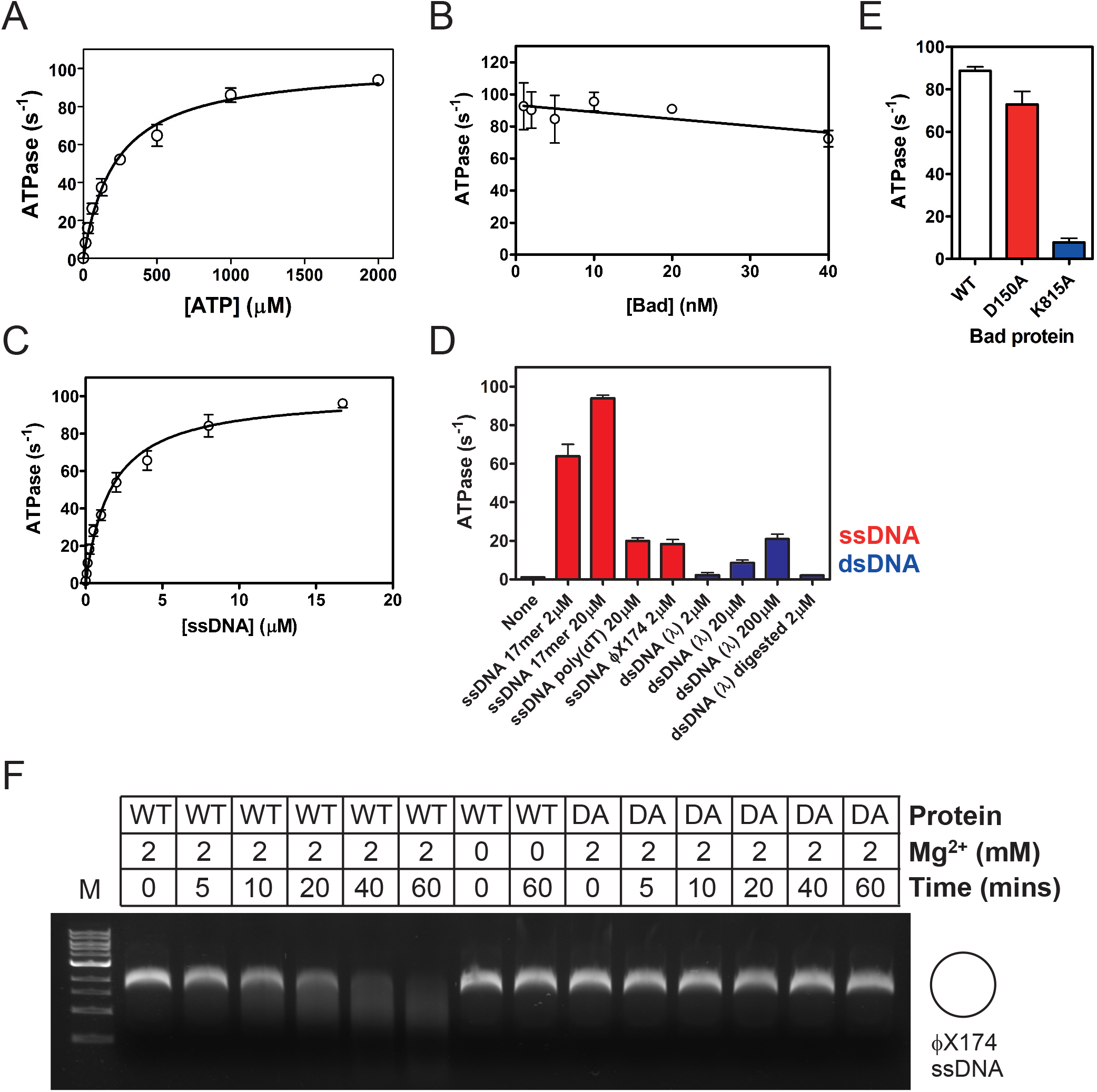
Bad is a ssDNA-dependent ATPase and ssDNA endonuclease. (A) Bad hydrolyses ATP with Michaelis-Menten kinetics (K_M_ = 223 ± 24 μM and k_cat_ = 101 ± 4 s^-1^). These measurements were made in the presence of saturating (20 μM) ssDNA oligonucleotide. (B) The ATP turnover rate as a function of Bad concentration provides no evidence for cooperativity in ATP hydrolysis. (C) Hydrolysis of ATP is ssDNA-dependent with half-maximal stimulation at K_DNA_ = 1.66 ± 0.22 μM. These measurements were made in the presence of saturating (2 mM) ATP. (D) ATPase activity was measured with a variety of different nucleic acid co-factors. All single-stranded DNA molecules are effective activators of the ATPase activity, whereas dsDNA molecules are very poor co-factors. The presence or absence of DNA ends does not dramatically affect the ATPase rate. (E) Comparison of the ATPase activity of wild type Bad, nuclease-dead mutant (D150A) and helicase-dead mutant (K815A) at saturating ATP and DNA concentrations. (F) Single-stranded DNA endonuclease activity was assessed in the absence of ATP by incubating Bad with circular ssDNA as described in the Methods. In the presence of Mg^2+^ ions, wild type Bad completely degraded the DNA substrate, whereas the Bad D150A (DA) mutant showed no activity under the same conditions.

### Bad is a 5’-to-3’ DNA translocase and helicase

To characterise the anticipated DNA motor activity of Bad, we first employed classical translocase and helicase assays to establish the existence and polarity of any such activity (27). We initially used the nuclease mutant (D150A) and “low free magnesium” conditions (see the **Methods**) in order to avoid complications associated with degradation of the DNA substrates. Single-stranded DNA translocation was monitored using a streptavidin displacement assay (20). In this assay, oligonucleotides are labelled with biotin at either the 5’ or the 3’ end, and then bound to streptavidin. Translocating motor proteins are typically able to displace the streptavidin in an ATP-dependent manner, but only if they translocate towards the target biotin moiety, and this can be monitored as the loss of a gel shift using native gel electrophoresis. The Bad D150A mutant protein was able to efficiently displace streptavidin from the ends of 3’-biotinylated oligonucleotides, and this activity was completely dependent on ATP and free biotin (**Figure 3A**). The free biotin acts to trap displaced streptavidin and prevent re-binding to the oligonucleotide. Therefore, the “no biotin” experiment serves as a control to show that the streptavidin has been displaced from the oligonucleotide rather than having been cleaved from the oligonucleotide by nuclease activity. In contrast, the Bad D150A protein showed no detectable streptavidin displacement activity on 5’-biotinylated oligonucleotides (**Figure 3B**). These data suggest that Bad is a 5’-to- 3’ ssDNA motor protein. DNA unwinding activity was monitored using classical strand displacement helicase assays for both wild type and mutant Bad proteins under low free magnesium conditions (22). These assays determine DNA unwinding polarity by comparing activity on three test substrates, two of which comprise short DNA duplexes flanked by either a 5’- or a 3’-ssDNA overhang, and one of which contains an equivalent duplex with no overhang. The Bad D150A protein was only able to efficiently unwind duplexes flanked by 5’-terminated ssDNA overhangs (**Figure 4**). This data is consistent with the translocase assays, shows that Bad displays 5’-to-3’ polarity, and classifies the enzyme as a SF1B helicase (28).

**Figure 3.**
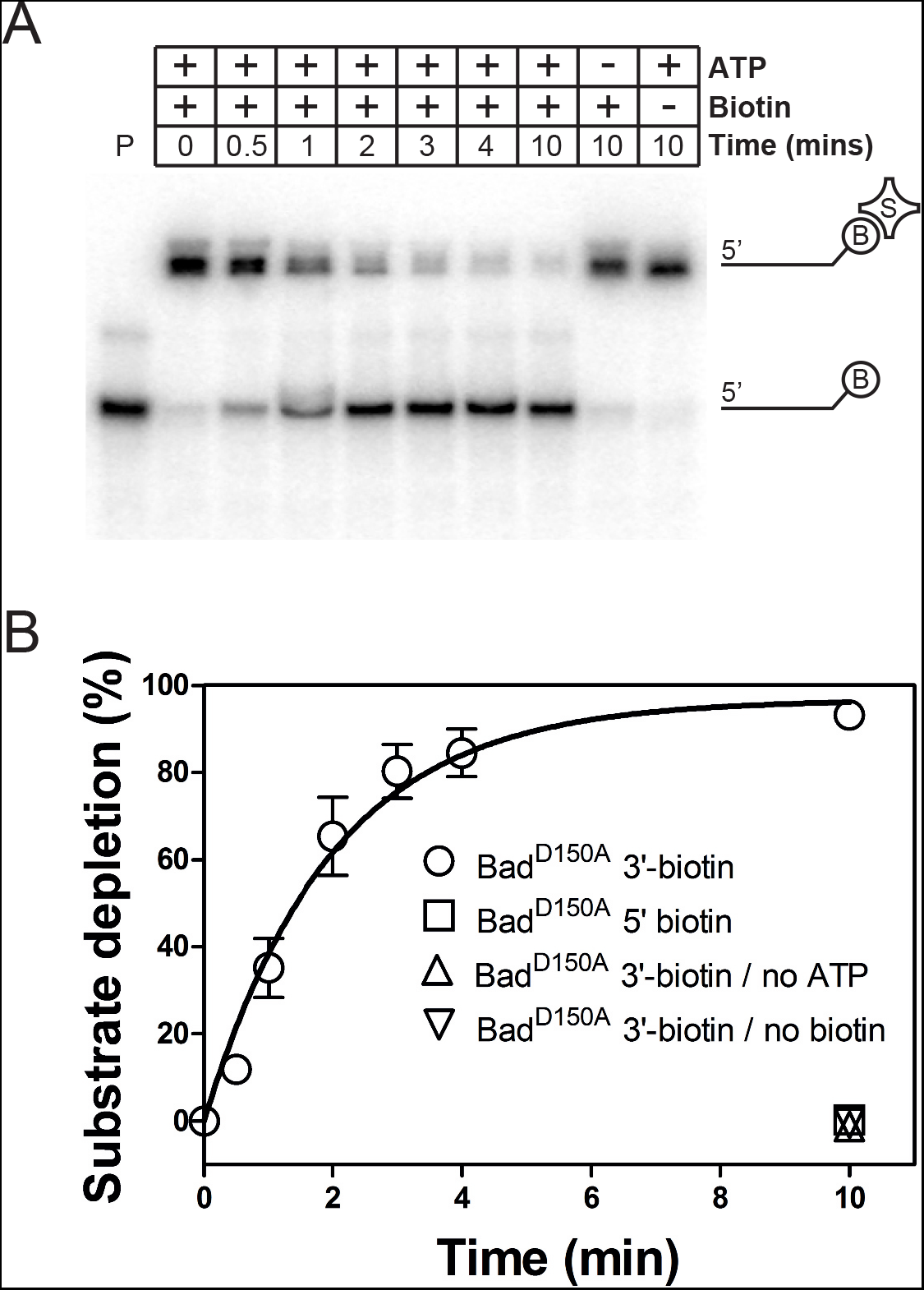
Bad is a 5′-to-3′ single-stranded DNA translocase. (A) Representative streptavidin displacement assay for Bad^D150A^ protein and an oligonucleotide labelled with biotin at the 3’ end. (B) Quantification of the gel-based assays to compare the streptavidin displacement kinetics on 3’- and 5’-biotin labelled DNA. Control experiments on 3’-labelled DNA in the absence of either ATP or biotin are also shown. Bad D150A shows no 3′-5′ translocase activity. The data for the 3’-labelled DNA in the presence of both ATP and biotin were well fit to a single exponential to yield an apparent rate constant k_obs_ = 0.48 min^-1^.

**Figure 4.**
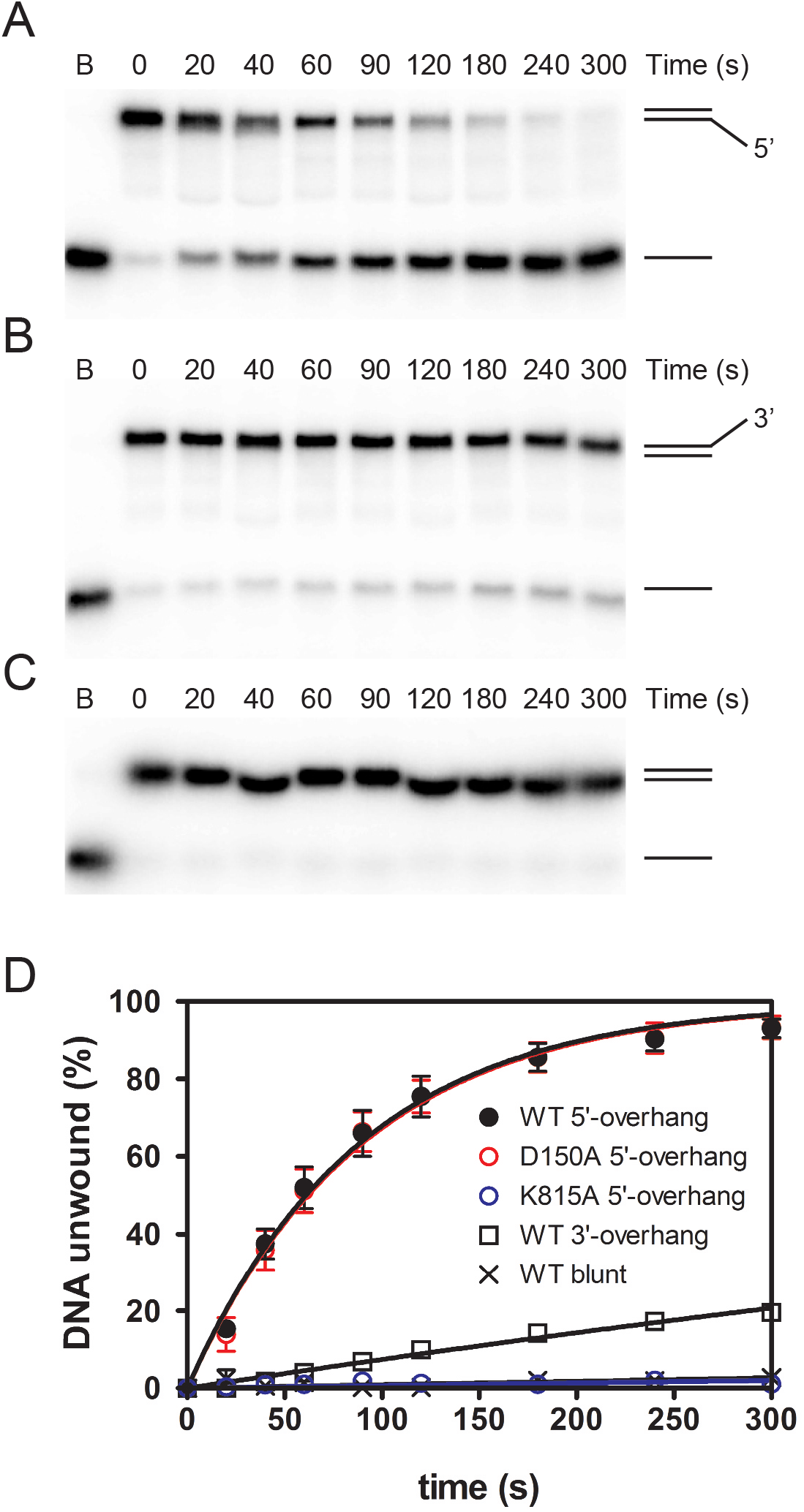
Bad is a 5′-to-3′ DNA helicase. (A-C) Representative helicase assay gels for wild type Bad protein acting on duplex DNA substrates containing either 5’-terminated, 3’-terminated, or no ssDNA overhang as indicated by the cartoons. (D) Quantification of helicase gels for wild type protein with each of the three DNA substrates performed in low free Mg^2+^ ion conditions (see Methods for details). Data for Bad^D150A^ and Bad^K815A^ are also shown for the 5’-terminated ssDNA overhang substrate only to allow comparison with wild type. Where appropriate the data were fit to a single exponential rise to yield apparent rate constants for unwinding (WT 5’-substrate = 0.66 min^-1^, D150A 5’-substrate = 0.66 min^-1^, WT 3’-substrate = 0.05 min^-1^).

### Bad displays coupled helicase and nuclease activities

We next analysed the activity of Bad under high free Mg^2+^ conditions (see the **Methods**) which promote both helicase and nuclease activity. This resulted in the formation of different and more complex unwinding products (**Figure 5**). For the junction with a 5’-terminated ssDNA overhang, wild type Bad both unwound and degraded the labelled DNA strand whereas the nuclease-dead mutant (D150A) only unwound it (**Figure 5A**). Interestingly, the helicase-dead mutant (K815A) produced a highly specific-cleavage product, suggesting that it binds to this 5’-overhang substrate in a preferred orientation that leads to precise endonucleolytic cleavage when ATP hydrolysis cannot take place. No such product was formed with this mutant protein on DNA molecules containing a 3’-ssDNA overhang or with blunt ends (**Figure 5B**). Using mass spectrometry, the position of this endonucleolytic cleavage event was mapped to a position on the 5’-overhang that was 13 nucleotides from the ss-ds junction (**Supplementary Figure 4**). Experiments with DNA junctions containing different duplex and 5’-overhang lengths showed that this cleavage position was always 13 nucleotides away from the ss-ds junction rather than being measured relative to the free 5’-end (**Supplementary Figure 4**). These data suggest that Bad specifically recognises the ss-dsDNA junction within a 5’-overhang substrate.

**Figure 5.**
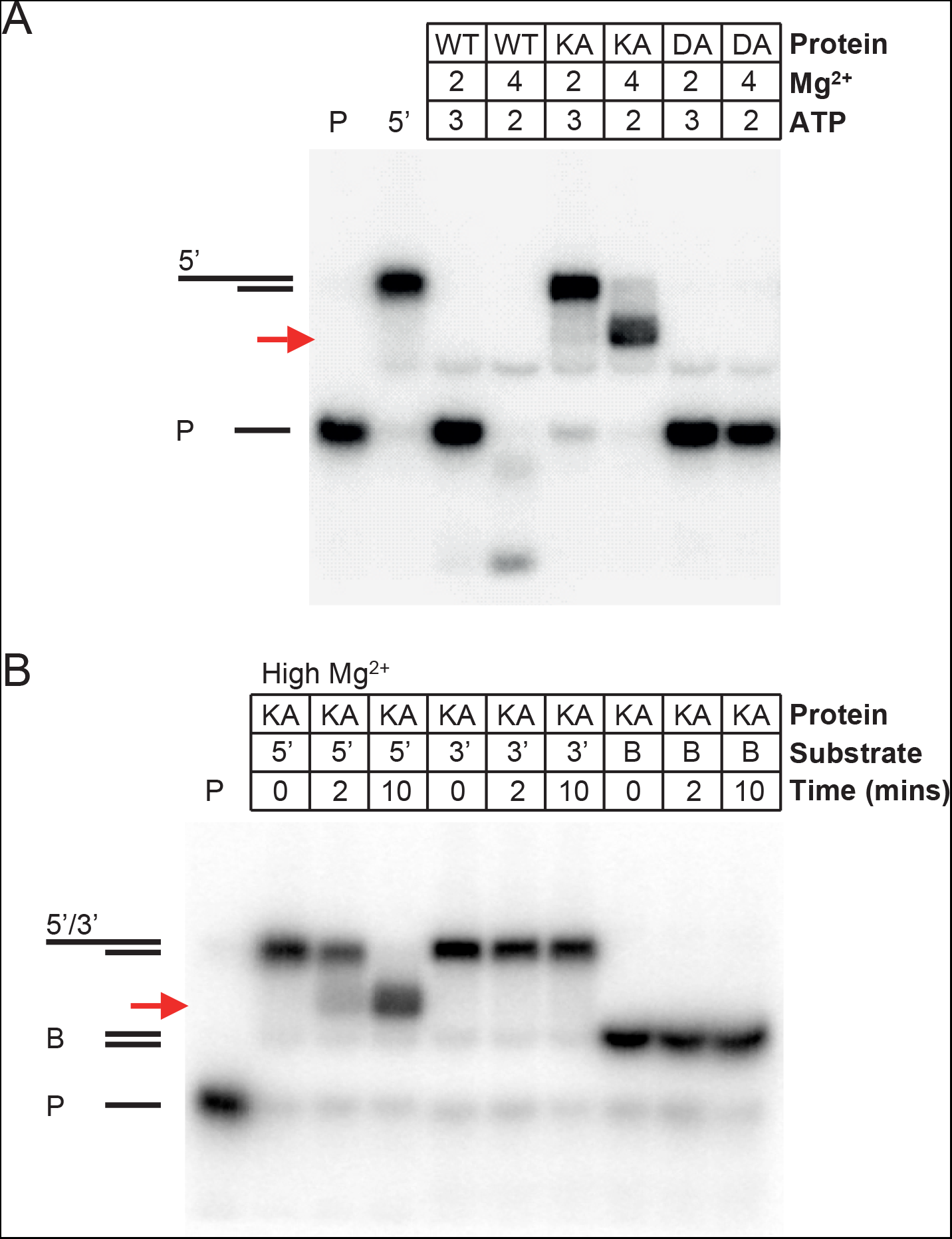
Bad displays coupled helicase and nuclease activities and binds in a well-defined orientation to 5’-overhang DNA substrates. (A) Helicase assays were performed with wild type or mutant Bad protein and 5’-terminated DNA substrates in either low or high free Mg^2+^ ion conditions. (B) Helicase assays were performed with the helicase-dead (K815A) mutant on 5’-terminated overhang, 3’-terminated overhang or blunt DNA substrates. In both panels, the red arrow indicates the position of an endonucleolytic cleavage product formed uniquely by the ATPase-dead mutant on the 5’-overhang substrate. Lanes containing the substrate only (5’) and the free short ssDNA product of unwinding (P) are labelled.

### Single molecule analysis of Bad reveals a fast and processive DNA motor

The ability of bulk helicase assays to provide mechanistic insight into DNA translocation and unwinding reactions is limited. This is because the observed activity is a measure of not only the DNA translocation and unwinding, but also of association/dissociation of the helicase and any failed unwinding events caused by lack of processivity or duplex re-annealing. These complications can be side-stepped by employing single molecule techniques in which translocation and unwinding are either directly observed or inferred from changes in the mechanical properties of the substrate DNA (29). Therefore, we used a magnetic tweezers (MT) approach to monitor the dynamics of DNA unwinding by Bad (**Figure 6**). We designed a ∼6.6kbp DNA substrate containing a free 5’-terminated poly-T ssDNA (37 nt) to act as a loading site for the enzyme (**Figure 6A**). We named this substrate MT1. DNA substrates were attached at one end to the surface of a flow cell and at the other end to paramagnetic beads. External magnets were used to apply force in order to extend and/or twist the DNA, while the Z height of the bead was monitored (**Figure 6B**). DNA unwinding can be monitored in this set-up because single- and double-stranded DNA display different force extension curves (30). In high applied-force (F≥6 pN) regimes, ssDNA is longer than duplex DNA and so helicase activity leads to an increase in the Z position of the bead (**Figure 6C**). Under low forces (F<6 pN), single-stranded DNA is shorter than duplex and unwinding leads to a reduction in the height of the bead (31, 32) (**Figure 6D**).

**Figure 6.**
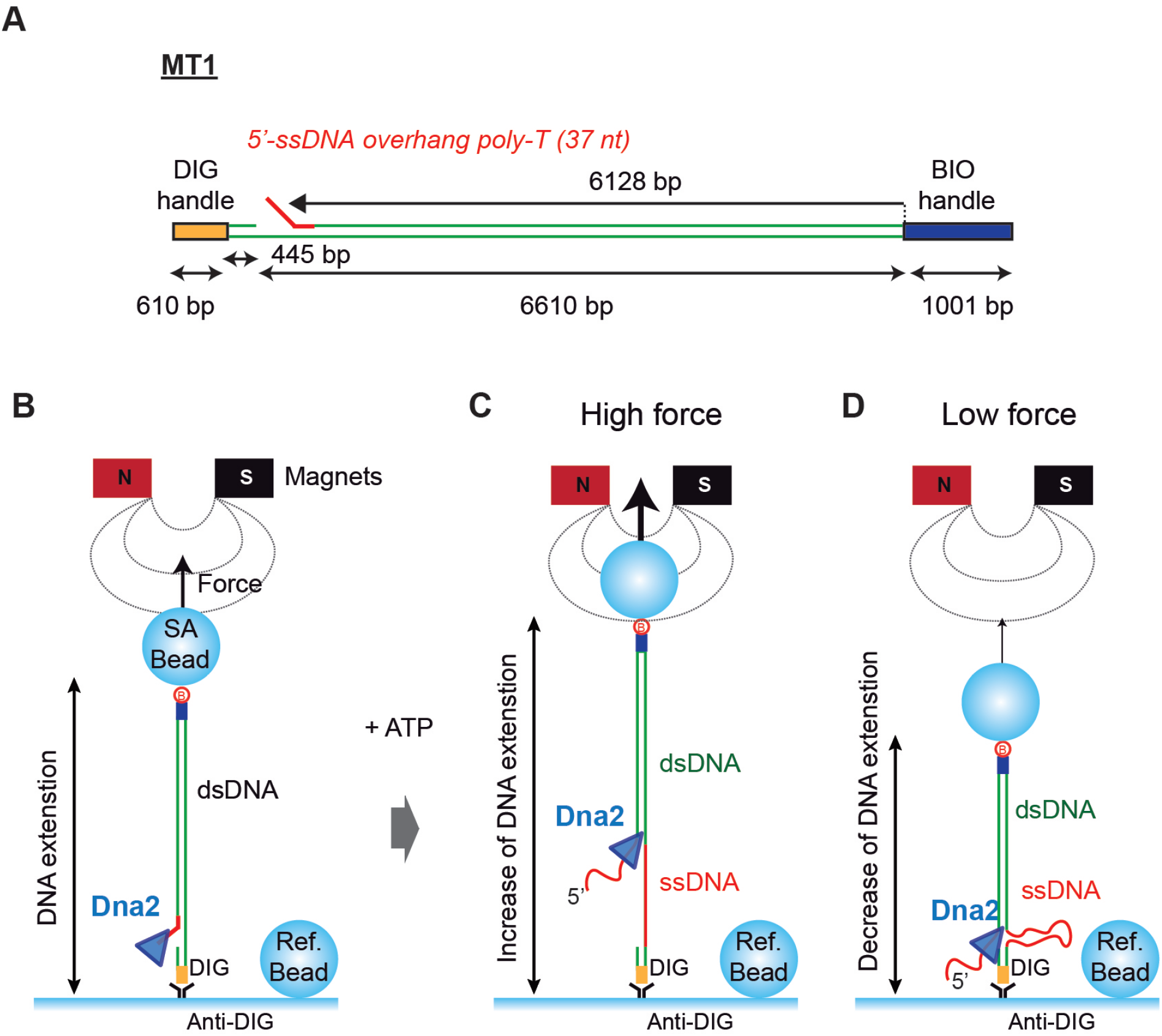
Magnetic Tweezers assay for monitoring DNA unwinding by Bad. (A) Schematic representation of the MT1 DNA substrate employed for the single molecule unwinding assay. It contains two handles labelled with biotins and digoxigenins which bind to the magnetic bead and glass surface, respectively. The insert (∼6.5 kbp) contains a 5’-ssDNA overhang at ∼6.1 kbp from the biotinylated DNA end. (B) In the experimental set-up, tethered DNA molecules are incubated with Bad proteins in the flow cell and Bad is loaded onto the 5’-ssDNA overhang. Note that, when a protein displays DNA unwinding (helicase) activity, the bead is expected to either (C) increase in height if the restraining force on the bead is high (because ssDNA is longer than dsDNA under these conditions) or (D) decrease in height under a low force regime (in which ssDNA is compacted compared to duplex).

In our initial experiments, it became apparent that neither an intact duplex, nor a duplex containing a site-specific nick were unwound by wild type or mutant Bad (**Supplementary Figure 5**). This was unsurprising, given that our bulk helicase assays had suggested that fully duplex DNA was a poor substrate, and that Bad bound to 5’-overhang substrates in a preferred orientation. Therefore, Tethered MT1-DNA-magnetic beads were incubated with Bad^D150A^ (the nuclease-dead mutant) at 8 pN applied force in the presence or absence of ATP. At all concentrations of Bad^D150A^ tested, ATP-dependent helicase activity was observed as many cycles of unwinding (U) and re-hybridization (R) in both high and low free Mg^2+^ion conditions (a representative trace is shown in **Figure 7A**). In contrast, unwinding by the wild type enzyme was only observed under conditions of low free Mg^2+^ ions and high ATP which suppress the nuclease activity (**Supplementary Figure 5**). This is consistent with the bulk data presented above and suggests that the nuclease activity of the wild type enzyme inhibits its own helicase activity, presumably either by efficiently cleaving the 5’-terminated loading strand from the substrate and/or by cutting the DNA track ahead of the SF1B motor domain. Additional control experiments without ATP or with the Bad^K815A^ (helicase-dead mutant) did not show any activity (**Supplementary Figure 5**). All of the further experiments described below use the nuclease mutant in low free Mg^2+^ conditions to minimise nuclease activity.

**Figure 7.**
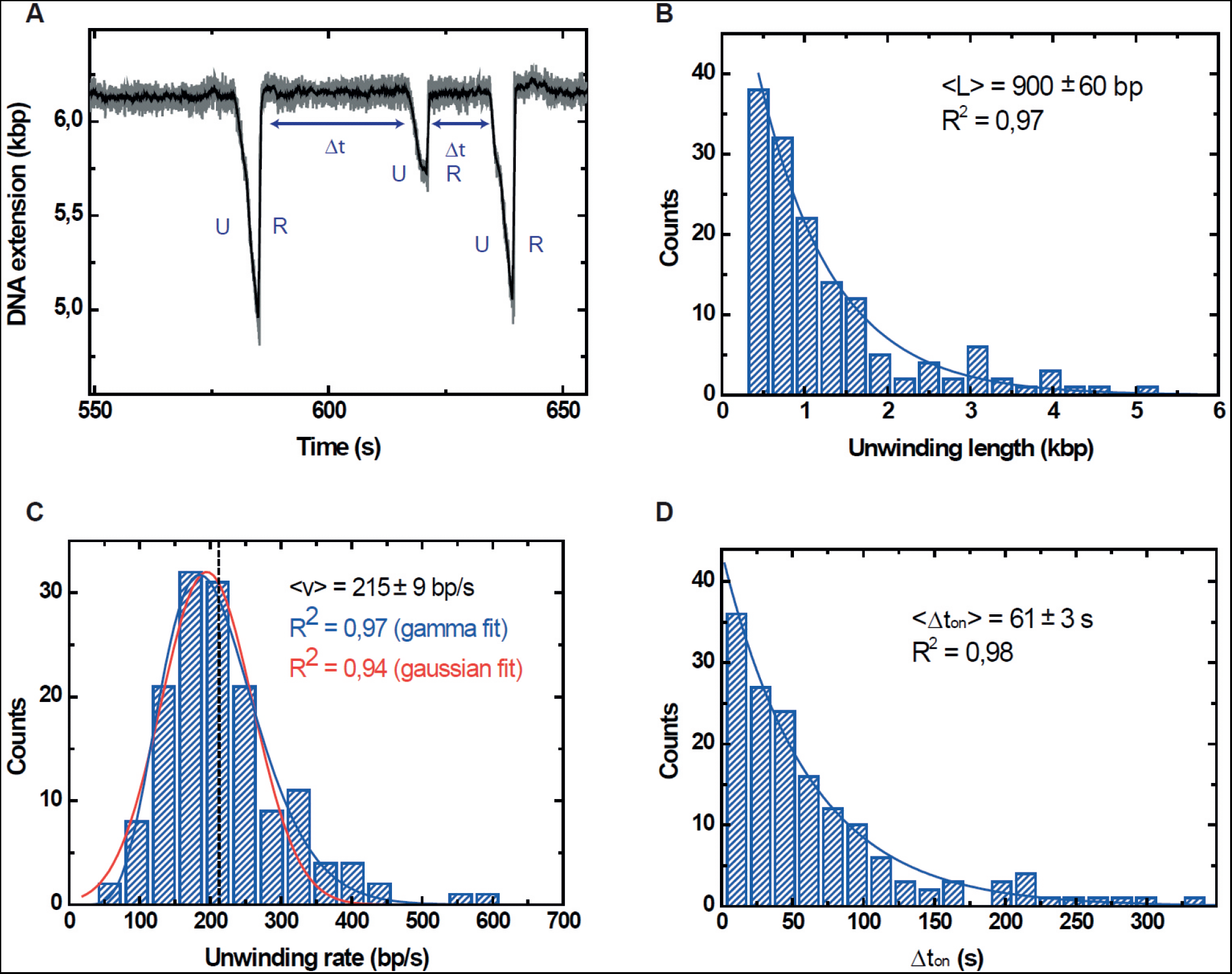
Dynamics of DNA unwinding by Bad at low protein concentration. (A) Typical unwinding trace measured at 30 nM Bad concentration and 8 pN force shows unwinding (U) and rapid rehybridization (R) events. We observe a decrease in DNA extension for the unwinding phase followed by a rapid increase in DNA extension when the enzyme detaches from DNA allowing a rehybridization and recovery of the initial DNA extension. (B) The distribution of the unwinding length events decays exponentially governed by the mean unwinding length event <L> = 900 ± 60 bp. (C) The distribution of the unwinding rate events was fitted with a Gamma function (blue) and with a Gaussian function (red). The quality of the Gamma and Gaussian fits are quite similar (see R^2^ values). The dashed line indicates the mean value of the rate. In total 57 tethered beads were analysed showing a total of 147 unwinding events. (D) Distribution of the dwell time between events Δ*t* decays exponentially governed by the mean time Δ*t* = 61 ± 3 *s*.

At a relatively low (30 nM) concentration of Bad, the observed unwinding length was exponentially-distributed, as is expected under standard models for helicase processivity (33), with an average distance travelled of 900 ± 60 bp (**Figure 7B**). The translocation rate distribution was well-fit to a Gaussian function with a mean value of 215 ± 9 bps^-1^ (**Figure 7C**). Finally, the dwell time between unwinding events (*Δt*) was exponentially-distributed with a time constant of 61 ± 3 s (**Figure 7D**). Pausing, backsliding and changes of rate during unwinding were rare under these conditions (see **Table 1** and **Supplementary Figure 6** for definition, quantification and examples of such events). We also characterised the DNA unwinding activity of Bad^D150A^ at a higher fixed concentration (160 nM) and similar unwinding and rehybridization events were observed (**Figure 8A**). However, although the frequency of backsliding remained similar, pauses during both unwinding and rehybridization now occurred more frequently (**Table 1**). From a total of 184 events, 21% and 37% of traces showed pauses in the unwinding and rehybridization respectively. The observed lengths of unwinding events were exponentially distributed with an average distance travelled of 1.18 kbp (**Figure 8B**). However, the rates of the unwinding events were now poorly described by a gaussian distribution. Instead, fitting to a gamma function showed an average value of 159 ± 6bps^-1^ (**Figure 8C**). The distribution of the dwell times (Δ*t*) between two unwinding events observed on the same DNA molecule was exponentially distributed with a shorter time constant of 12 s (**Figure 8D**).

**Figure 8.**
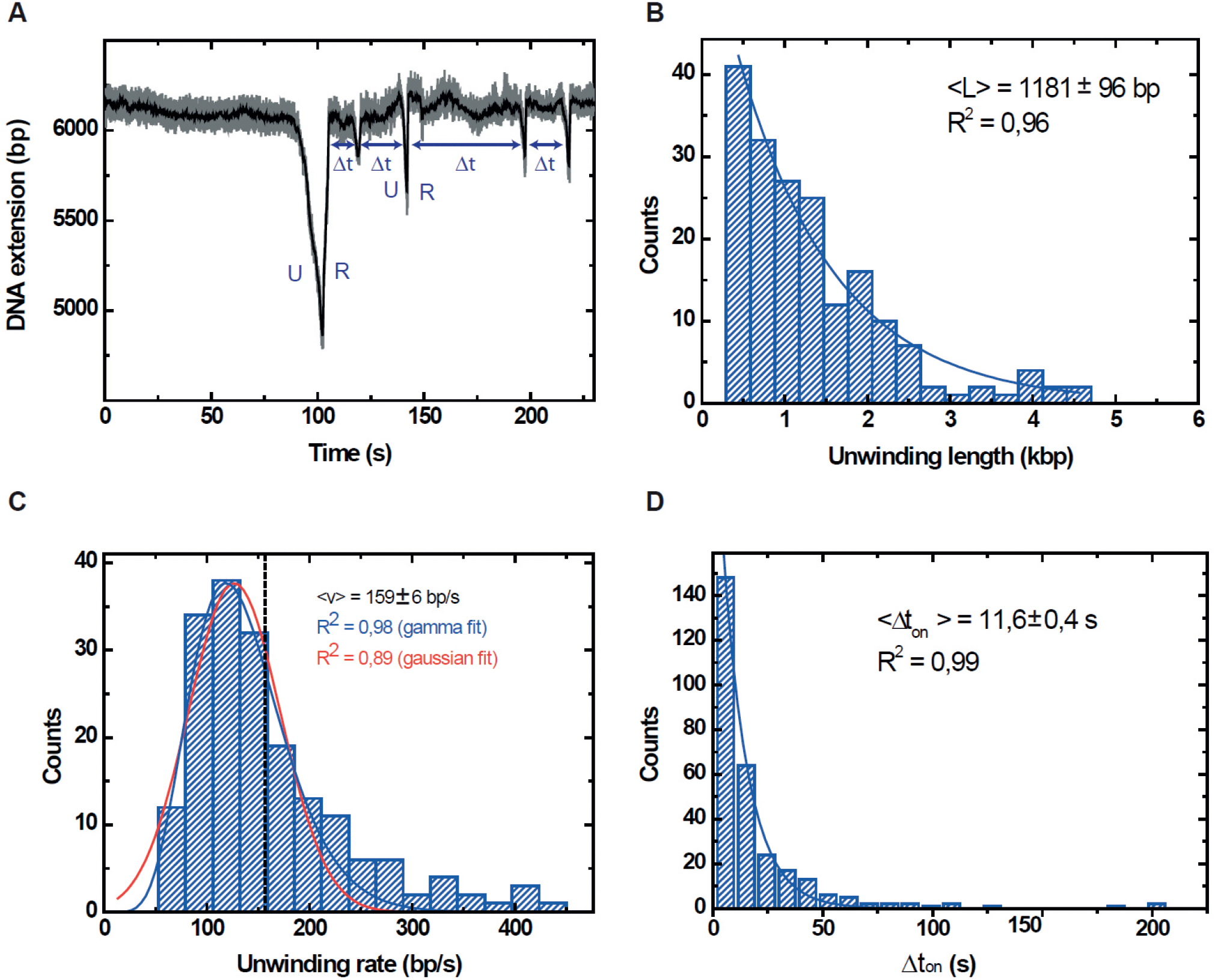
Dynamics of DNA unwinding by Bad at high protein concentration. (A) Typical unwinding curve at 163 nM Bad and 8 pN force shows several unwinding (U) and quick rehybridization (R) events. (B) Distribution of the unwinding length events decays exponentially governed by the mean unwinding length event <L> = 1181 ± 96 bp. (C) The distribution of the unwinding rates was fitted with a Gamma function (blue) and with a Gaussian function (red). The dashed line indicates the mean value of the rate. In total, 184 unwinding events from 92 tethered beads were analysed. (D) Distribution of the dwell time between events Δ*t* decays exponentially governed by the mean time Δ*t* = 11.6 ± 0.4 *s*.

**Table 1.**
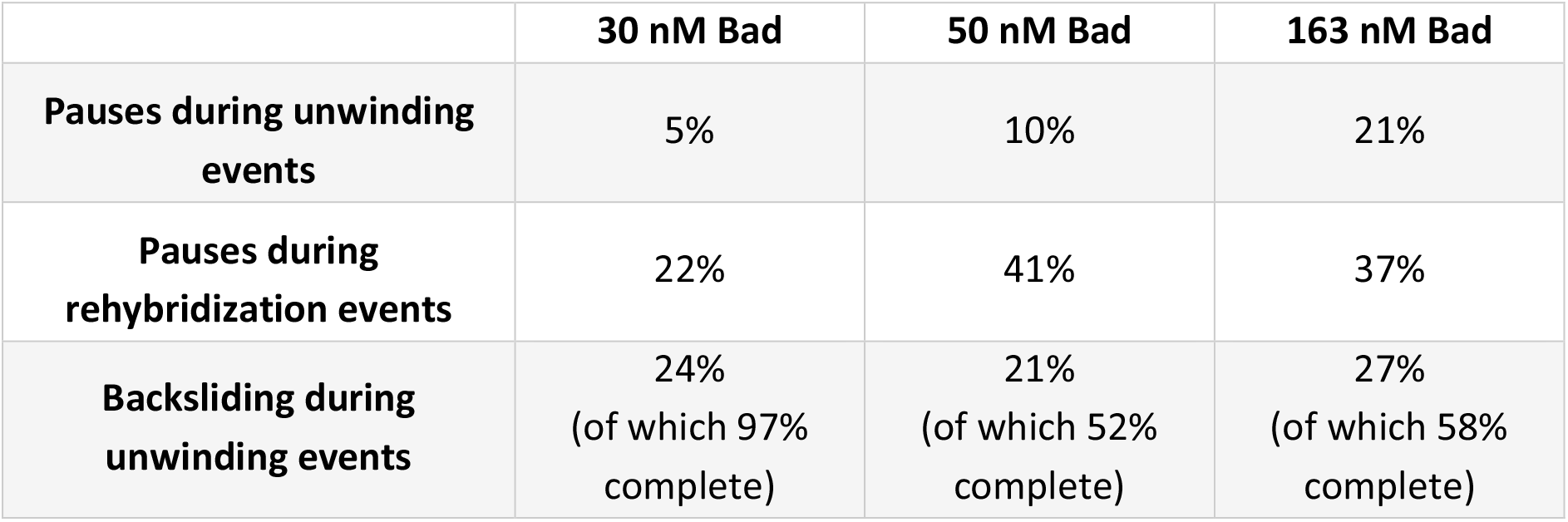
Frequency of pauses and backsliding during DNA unwinding and rehybridization. Higher Bad concentrations result in more pauses during unwinding and rehybridization. The frequency with which backsliding events are observed is approximately constant, but a greater proportion of backsliding events lead to complete rehybridization if the Bad concentration is low.

In summary, we found that the apparent translocation rate decreases at higher Bad concentrations, whereas the initiation frequency, the pause frequency and processivity all increase (**Figure 9A and Table 1**). These observations can all be explained by the idea that many more initiation events occur at high concentrations of Bad, such that multiple Bad molecules may be translocating on a single substrate at the same time. This could hinder DNA translocation, leading to pausing, spontaneous changes in the rate of unwinding and a complex rate distribution as observed, but might also improve the processivity by disfavouring re-hybridisation. The constant frequency of the backslide events regardless of protein concentration suggests that these are an intrinsic property of the functional form of the enzyme and are the result of re-initiation of unwinding by the same enzyme, rather than an artefact caused by overloading of the DNA substrate. We hypothesize that backsliding results from dissociation of the DNA motor domains of Bad from the DNA track, but that the protein can retain a loose grip on the substrate (probably with the non-translocated strand for reasons discussed below), allowing it to re-engage and resume translocation. The relationship between the dwell time Δ*t_on_* and [Bad] suggests that initiation and re-initiation events from the loading site are caused by binding of Bad from free solution and allow us to calculate the second order rate constant for this process (defined as *k_on_* = 1 / (< Δ*t_on_* >* [*Bad*])) as *k_on_* = 5.9×10^5^ M^-1^ s^-1^ (**Figure 9B**).

**Figure 9.**
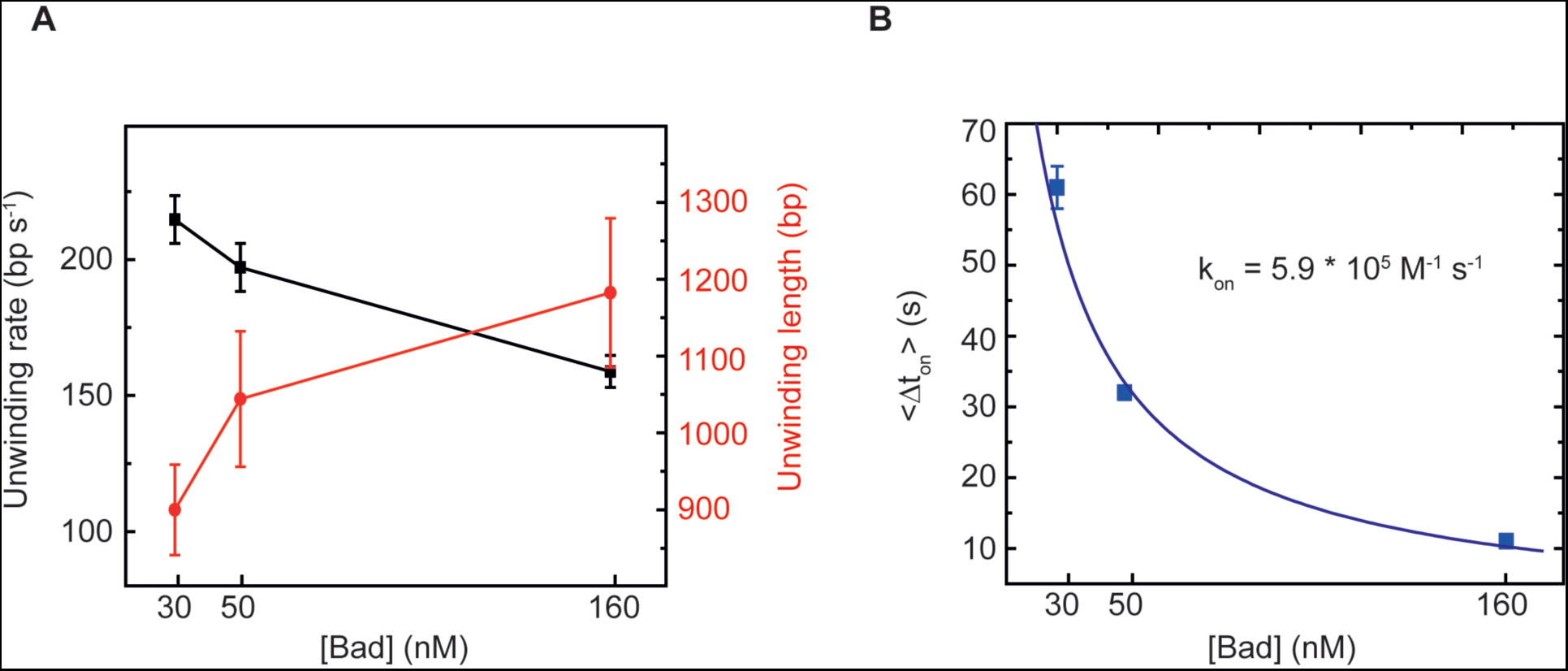
Effect of Bad concentration on DNA unwinding parameters. (A) Unwinding rate and unwinding length at different protein concentrations (30 nM, 50 nM, 163 nM). The mean value and standard error of the mean are shown for different protein concentrations (30 nM, 50 nM, 163 nM). (B) Plot of the mean dwell time < Δ*t* > *versus* Bad protein concentrations. Data were fitted with a simple hyperbolic function to obtain the binding rate constant (see main text).

### DNA unwinding by Bad is coupled to single-stranded DNA loop extrusion

An interesting and somewhat unexpected feature of these traces is that the activity of Bad manifests itself as an ATP-dependent *decrease* in the bead height despite the high (8 pN) restraining force. At this applied force, the ssDNA product is actually expected to be longer than the duplex substrate (30) (**Figure 6B**). Indeed, even at a restraining force of 14 pN, we found that Bad caused the bead height to processively *decrease*, and the traces were very similar to those measured at 8 pN (data not shown). To confirm that we were indeed observing DNA strand separation, we also performed experiments in the presence of bacterial single-stranded DNA binding (SSB) protein. We reasoned that, if ssDNA is formed during ATP-dependent translocation, then DNA rehybridization events (marked R) should be much slower in the presence of SSB. In these experiments, tethered DNA molecules were incubated with both Bad and SSB proteins at 8 pN applied force (**Supplementary Figure 7**). Under these conditions, although rehybridization did still occur, the overwhelming majority (93%) of the events showed a dramatically slower rehybridization. Moreover, rehybridization could be completely eliminated at higher [SSB]. Interestingly, “*backslide*” events were also substantially reduced (to ∼10% of the total events analyzed) in the presence of SSB.

We considered several possible models for how Bad activity might decrease the height of the bead. Firstly, the bead height change could be caused by an experimental artefact, such as the Bad protein sticking to the flow cell surface during translocation. This is unlikely because the observed enzyme activity requires the 5’-ssDNA loading site, this is located at a position distant from the surface, and the traces provide no evidence to suggest that binding of Bad causing the loading site to interact with the surface. This is true even when the loading site is re-positioned much further from the surface (data not shown). Therefore, we favour an alternative possibility in which the Bad monomer contains multiple DNA binding sites and remains bound near the loading region of the substrate while translocating on 5’-strand (**Figure 10**). In this scenario, movement along the DNA would cause looping on the non-translocating strand, leading to the formation of a ssDNA loop and the observed decrease in bead height.

**Figure 10.**
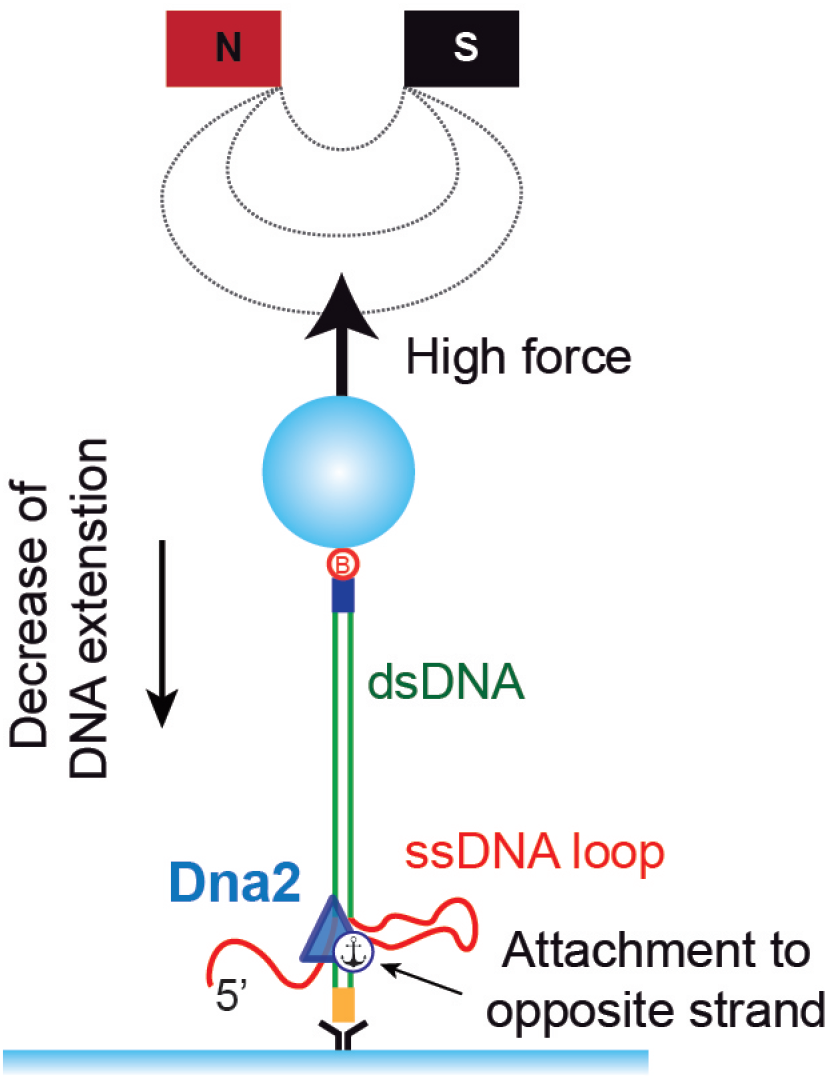
Model for Bad-dependent single-stranded DNA loop extrusion. The decrease in bead height we observe due to ATP-dependent Bad translocation and unwinding can be explained by a model in which the non-translocated (opposite) strand forms a ssDNA loop that is extruded from the enzyme complex during movement. If the enzyme does *not* retain a contact with the DNA behind the flap on the translocated strand, then there is no topological constraint to movement and the decrease in bead height will simply be equivalent to the distance moved forward by the enzyme. This idea can be tested using nicked DNA substrates.

### The arrest of Bad at single-strand nicks confirms the loop extrusion model

In a model where Bad remains bound to parts of the DNA substrate other than the translocating strand, the reduction in bead height associated with DNA translocation may either be explained by simple loop extrusion as we propose, or by the introduction of positive writhe in the DNA ahead of the translocating motor. In the first scenario, the decrease in bead height would be directly equivalent to the distance travelled into duplex DNA by Bad. In the second scenario, the interpretation of the relationship between translocation rate (in base pairs) and observed bead height (in microns) would be complex, with relatively small translocation events causing larger effects on the beads. To formally discriminate between such models and to test how Bad translocation was affected by damage to either strand of the duplex, we next performed experiments with substrates containing nicks. The substrate DNA-nick-top contains a nick in the 5’-to-3’ translocated strand, whereas DNA-nick-bottom is nicked in the 3’-to-5’ non-translocated strand (**Figure 11A**). We reasoned that if translocation ceased at a distance equivalent to the distance between the loading site and the lesion, then we were observing simple loop extrusion. Initial control experiments using nicked DNA molecules without a 5’-ssDNA overhang as a loading site showed no activity, confirming that nicks do not themselves act as productive loading sites for Bad (**Supplementary Figure 5A**). Experiments performed at 30 nM Bad using a substrate with a nick in the top strand (**Figure 11A**) revealed that the bead never moves further than the distance between the loading site and the nick, and the length distribution is Gaussian-distributed suggesting that translocation is prematurely arrested at the approximate position of the nick (**Figure 11B**). In complete contrast, experiments using a nick in the bottom strand showed similar unwinding length distributions to the MT1 control substrate (**Figure 11A**), being well-fitted by an exponential function and giving a mean value of 870 ± 117 bp (**Figure 11B**). Together, these data suggest that nicks on the translocating strand strongly inhibit Bad translocation, confirm the 5’-3’ polarity of Bad measured in bulk assays, and strongly suggest that DNA translocation and unwinding are accompanied by simple loop extrusion on the non-translocated strand. This mode of unwinding is also consistent with our observation that the bead is not released when Bad translocation proceeds past the position of the nick on the DNA-nick-bottom substrate.

**Figure 11.**
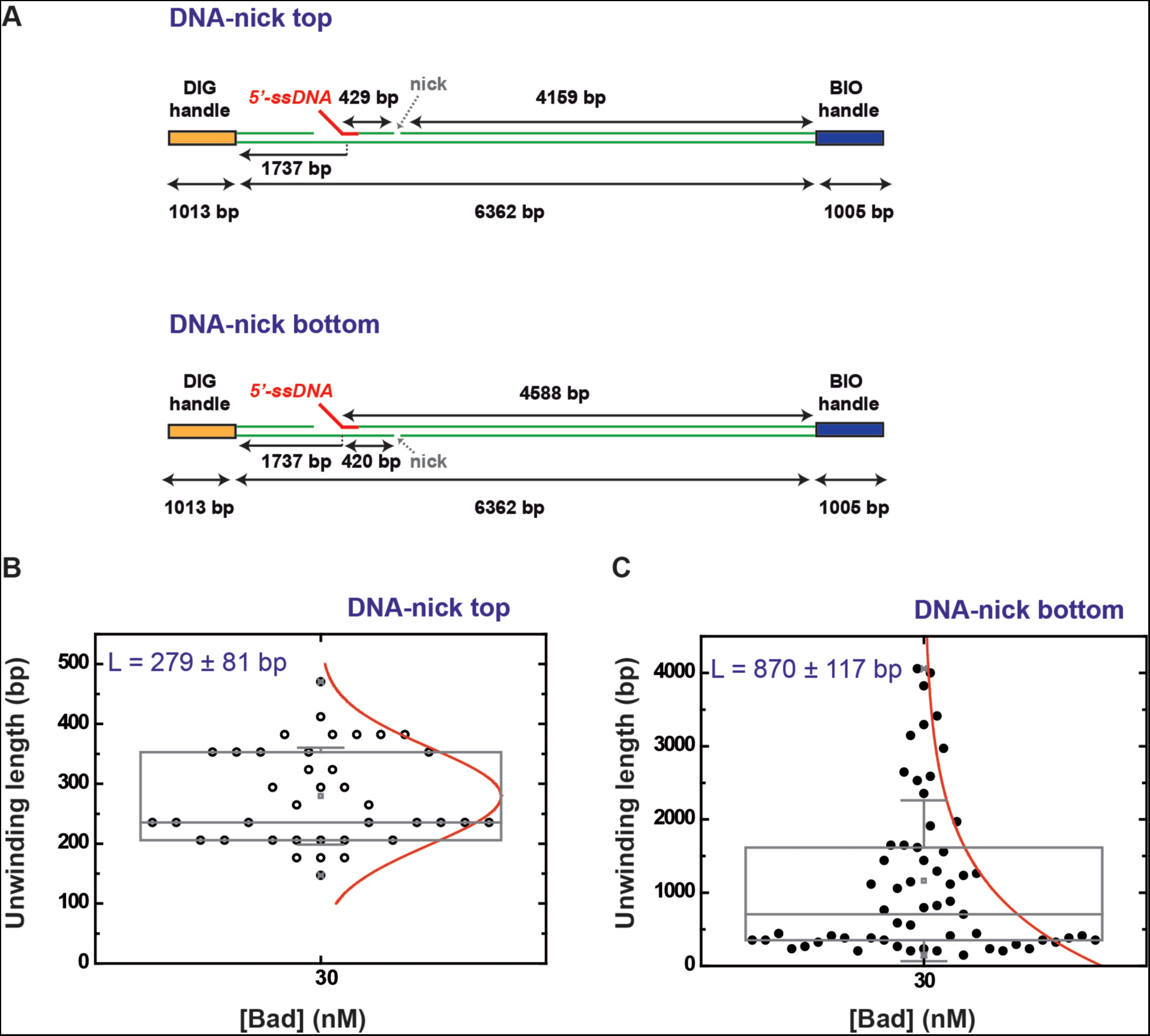
Unwinding activities using nicked DNA substrates. (A) Structure of the substrates used. (B) Box plot graphs of the observed unwinding length for DNA with a nick in the top (translocating) strand and for (C) DNA with a nick in the bottom strand. Experiments were performed at 30 nM Bad concentration with the nuclease mutant. Box plots indicate the median, 25th, and 75th percentiles of the distributions and the whiskers show the standard deviation. For both data sets, the red line shows the distribution of the data which is gaussian for the top strand nick but exponential for the bottom strand nick.

## DISCUSSION

In this work we identified and characterised a bacterial helicase-nuclease fusion with primary structure homology to the eukaryotic DNA replication and repair factor DNA2. This enzyme, which we call Bad, is rare and sporadically distributed in the bacterial domain. This find was unexpected given that DNA2 had been considered exclusively eukaryotic in origin (2). We showed here that the biochemical behaviour of Bad is highly similar to that of eukaryotic DNA2 proteins in many respects. Both Bad and DNA2 contain an Fe-S cluster that is important for structural integrity and activity (9). Moreover, they both possess ATPase activity, 5’-3’ ssDNA motor activity, DNA unwinding activity and nuclease activity *in vitro* (34, 35), and they both display a preference for binding and/or unwinding substrates with a 5’-flap (36, 37). Interestingly, the helicase activity of Bad is autoinhibited by its own nuclease activity (at least *in vitro*) as has also been shown for DNA2. This is presumably because the enzymes cleave DNA ahead of themselves; a counterintuitive activity that could suggest that they act in complex with other proteins (which might overcome this inhibitory effect by providing additional DNA binding sites), or that they are specifically designed to cleave 5’-flap structures until they are sufficiently short to prevent binding. Single molecule analysis revealed that Bad is a fast and highly processive DNA motor protein, and that DNA unwinding proceeds by the formation of a ssDNA loop on the non-translocated strand. In our magnetic tweezers set-up, this enables the enzyme to decrease the apparent DNA extension even against high restraining forces. Loop extrusion is an emerging feature of highly-processive DNA helicases and might assist DNA unwinding by dis-favouring re-annealling even in the presence of single-stranded DNA binding proteins (see (38) for discussion). The bacterial Bad system may provide an interesting model system for studying this activity, particularly using single molecule approaches. Previous analysis of yeast and human Dna2/DNA2 using magnetic tweezers did not provide evidence for ssDNA loop extrusion (12, 14). In experiments performed at ∼25 pN, DNA unwinding by Dna2 led to a progressive extension in apparent DNA length as expected for a canonical unwinding activity. This DNA strand separation was also found to be dependent on a single-stranded DNA binding protein (RPA), which was not the case here with Bad.

The physiological role of bacterial Bad proteins is unknown, but they are unlikely to be straightforward orthologues of eukaryotic DNA2 for at several reasons. Firstly, the Bad protein is not ubiquitous in bacterial cells, whereas eukaryotic DNA2 is ubiquitous and essential. Furthermore, the major roles played by DNA2 in eukaryotic organisms are apparently already provided by other enzymes in bacteria. For example, the AddAB- and RecBCD-type helicase-nucleases are responsible for DNA end resection to promote homologous recombination (15), and Okazaki fragments are processed by RNaseH (39), although the latter process remains surprisingly poorly understood in bacteria (40). Finally, Bad proteins all contain a central conserved domain that is not present in any eukaryotic DNA2 protein (**Supplementary Figures 1-3**). The structure and function of this domain is completely unknown, as it bears no primary structure homology to anything in the available databases other than uncharacterised Bad proteins. Moreover, domain prediction algorithms also fail to find remote homology with known structures (data not shown). One possible clue as to the cellular role of Bad can be found in the genome organisation of bacteria encoding this protein. In the *Geobacilli*, in all instances we investigated, the *bad* gene was always found neighbouring a predicted DNA methyltransferase related to the M subunit of TypeIII restriction enzymes (41, 42). Whole genome sequencing of *Geobacillus stearothermophilus* 10 (the organism from which the Bad protein studied here originates) using SMRT sequencing suggests that this enzyme methylates the N6-position of adenine in the sequence 5’-GCCAT-3’ (41). Therefore, Bad may be a component of a novel restriction enzyme or any other system which is regulated by DNA methylation. Note however that conventional TypeIII restriction-modification systems do not possess DNA motor activity and are instead ATP-dependent DNA sliding proteins (43). Moreover, even though the TypeI restriction-modification systems do contain *bona fide* DNA motor subunits, these are formed by Superfamily II “translocase” enzymes which move along DNA without unwinding. Therefore, it is plausible that Bad is a subunit of a novel class of restriction enzyme, which might unwind and degrade DNA concomitantly. This hypothesis will be the subject of future work.

## FUNDING

This work was supported by the Wellcome Trust (100401/Z/12/Z to MD). F.M.-H. acknowledges support from European Research Council (ERC) under the European Union Horizon 2020 research and innovation (grant agreement No 681299). The work in the Herrero-Moreno laboratory was also supported by a Spanish Ministry of Economy and Competitiveness grant, BFU2017-83794-P (AEI/FEDER, UE) to F.M.-H., and Comunidad de Madrid grants, Tec4Bio – P2018/NMT-443 and NanoBioCancer – Y2018/BIO-4747 to F.M.-H.

## ACKNOWLEDGEMENTS

We thank Prof. Mark Szczelkun for stimulating discussions regarding the mechanism of action of restriction enzymes that were relevant to this work and its interpretation. We thank Prof. Ralf Seidel for the kind gift of the pNLrep plasmid used for the production of our magnetic tweezers DNA substrates.

