## Supplementary Information for "Bulk and single-molecule analysis of a novel DNA2-like helicase-nuclease reveals a single-stranded DNA looping motor"

**Supplementary Methods**

Oligonucleotides used in ATPase, translocase and helicase assays.

| Name | Sequence |
|---|---|
| ODN1 | CAAGCGGGCGCCTCCCG |
| ODN2 | G(biodT)ACGTATTCAAGATACCTCGTACTCTGTACTGACTG ATCCTA |
| ODN3 | GTACGTATTCAAGATACCTCGTACTCTGTACTGACTCGGATCC (biodT)A |
| ODN4 | AACGCGCGGGGAGAGGCGGTTTGCGTATTGGGCGCTCTTCCG CTTCCTCGCTCACTGACT |
| ODN5 | CAATACGCAAACCGCCTCTCCCCGCGCGTT |
| ODN6 | AGTCAGTGAGCGAGGAAGCGGAAGAGCGCCCAATACGCAAAC CGCCTCTCCCCGCGCGTT |
| ODN7 | AACGCGCGGGGAGAGGCGGTTTGCGTATTG |
| ODN8 | ACTTATCGGTAGTCAGTGAGCGAGGAAGCGGAAGAGCGCCCAATACGCAAAC CGCCTCTCCCCGCGCGTT |

| Assay | Substrate | ODNs |
|---|---|---|
| ATPase | ssDNA 17mer | 1 |
| Translocation | 5′-3′ | 2 |
| | 3′-5′ | 3 |
| Helicase | 3′-5′ (60) | 4+5 |
| | blunt | 5+7 |
| | 5′-3′ (60) | 6+7 |
| | 5′-3′ (70) | 7+8 |

**Supplementary Figure Legends**

**Supplementary Figure 1** *Domain structure of bacterial DNA2-like (Bad) proteins.* (Top) Multiple sequence alignment of bacterial and archaeal DNA2-like proteins shown in WebLogo format (1). The bars to the left of the logo approximately delineate the overall architecture showing the N-terminal Fe-S associated RecB-family nuclease domain (red), a central domain (green) and a C-terminal Superfamily 1B helicase domain (blue). Aside from a short region of predicted coiled-coil, no information can be gleaned from bioinformatics approaches about the likely structure or function of the central green domain. Amino acid motifs that are characteristic of the Fe-S RecB-family nuclease are highlighted with red bars above the logo and include the flanking cysteine residues, nuclease motifs 1-3, and the RecB-family specific motif (2). Amino acid motifs 1-6 that are characteristic of SF1B helicases are highlighted with blue bars above the logo (3). The multiple sequence alignment was created by BLAST-based homology searching using a representative proteome database (4), followed by manual editing of the resulting alignment in COBALT (5) to remove sequences of significantly different length or those which did not conform to the three domain architecture indicated above. The sequences used for the final multiple sequence alignment were the Bad proteins from the following bacterial and archaeal species: *Geobacillus thermoleovorans* (WP_011230911.1), *Parageobacillus toebii* (WP_062677896.1), *Anoxybacillus flavithermus* (WK1 ACJ33552.1), *Methanohalobium evestigatum* (WP_013193644.1), *Methanosphaerula palustris* (WP_012618149.1), *Methanospirillum hungatei* (WP_011448513.1), *Methanoculleus marisnigri* (WP_011844558.1), *Methanoculleus marisnigri* (KUL02919.1), *Methanoculleus bourgensis* (WP_014866635.1), *Methanofollis liminatans* (WP_004038903.1), *Candidatus Competibacter denitrificans* (WP_048677101.1). (Bottom) Modelled structures of the N-terminal Fe-S nuclease (red) and C-terminal helicase (blue) domains of *G. stearothermophilus* Bad. The models were generated using Phyre2 and are based on homology to *B. subtilis* AddB and mouse DNA2 respectively.

**Supplementary Figure 2** *Domain structure of DNA2 from human and related species.* Multiple sequence alignment of higher eukaryotic DNA2 proteins shown in WebLogo format (1). The bars to the left of the logo approximately delineate the overall architecture showing the N-terminal Fe-S associated RecB-family nuclease domain (red) and a C-terminal Superfamily 1B helicase domain (blue). Amino acid motifs that are characteristic of the nuclease and helicase domains are labelled as in **Supplementary Figure 1**. The multiple sequence alignment was created by BLAST-based homology searching using mouse DNA2 in a representative proteome database (4), followed by manual editing of the resulting alignment in COBALT (5) to remove sequences of significantly different length or those which did not conform to the domain architecture indicated above.

**Supplementary Figure 3** *Domain structure of yeast Dna2 proteins.* Multiple sequence alignment of *S. cerevisiae* and closely-related DNA2 proteins shown in WebLogo format (1). The bars to the left of the logo approximately delineate the overall architecture showing the N-terminal yeast specific domain (purple), the Fe-S associated RecB-family nuclease domain (red) and a C-terminal Superfamily 1B helicase domain (blue). Amino acid motifs that are characteristic of the nuclease and helicase domains are labelled as in **Supplementary Figure 1**. The multiple sequence alignment was created by BLAST-based homology searching using *S. cerevisiae* Dna2 in a representative proteome database (4), followed by manual editing of the resulting alignment in COBALT (5) to remove sequences of significantly different length or those which did not conform to the domain architecture indicated above.

**Supplementary Figure 4** *Bad binds and cleaves a 5'-ssDNA overhang at a precise location relative to the ss-dsDNA junction in an ATP-dependent manner.* (A) Coupled helicase-nuclease assays were performed with wild type or mutant Bad protein in high free $Mg^{2+}$ ion conditions and with or without ATP. In order to distinguish between cleavage of DNA relative to the 5'-end of the long strand or to the ss-dsDNA junction two DNA substrates were used with different length top strands of 60 or 70 ntds. P is a marker DNA for the position of the unwound short DNA strand. (B) Schematic showing the Bad cleavage position on the 5'-60 DNA as revealed by mass spectrometric analysis of the products. The analysis revealed oligonucleotides with the exact mass of the short ssDNA strand (9384.1 Da), as well as the long DNA strand that had been cut precisely at a position 13 nucleotides from the junction to leave a 5'-phosphate (red arrow; 13183.5 Da). The 5'-portion of the long DNA strand was not detected in this analysis suggesting it may have been progressively degraded to the 13 ntd position (data not shown).

**Supplementary Figure 5. Single molecule control experiments.**

(A) Activity of nuclease-dead Bad mutant on torsionally-constrained (N = 2) and nicked (N = 4) DNA molecules. No unwinding events were observed. (B) Wild type Bad activity at high or low free $Mg^{2+}$ conditions as indicated. Unwinding events were only observed under low free $Mg^{2+}$ conditions in which the nuclease activity is suppressed. (C) Helicase-dead Bad mutant under low $Mg^{2+}$ conditions. N = 13, no unwinding events were observed. (D) Nuclease-dead Bad mutant activity on DNA containing a gap (25 bp). N = 34, 3 short unwinding and 1 backslide events were detected.

**Supplementary Figure 6. Examples of pauses, velocity changes and backsliding during Bad translocation.**

(A) Pauses during unwinding. (B) Pauses during rehybridization. (C) Velocity changes during unwinding. (D) Backsliding (rapid rehybridization following by restart of unwinding). Experiments were performed with 163 nM Bad under the low free magnesium ion conditions. Quantification of these events for a range of Bad concentrations is shown in the main text **Table 1**.

**Supplementary Figure 7. DNA unwinding and rehybridization in the presence of SSB.**

Examples of unwinding and rehybridization events are shown in (A) and (B) for the nuclease-dead mutant in the presence of 3 µM SSB. It was observed that rehybrization is always much slower than in the absence of SSB. Distribution of the (C) unwinding rate and (D) rehybridization rate events. Both were well fitted with a Gaussian function (blue lines). (E) Distribution of the unwinding length events decays exponentially governed by the mean unwinding length <L> = 275 ± 22 nm. The mean unwinding length in the presence of SSB is lower than that obtained without SSB proteins. Interestingly, the percentage of backsliding events in the data analyzed was reduced (10 %) in the presence of SSB. A total of 36 tethered beads were analysed showing a total of 131 unwinding events. (F) At 4 µM SSB, complete rehybridization events were never observed, although partial and short rehybridization events did still occur.

Supplementary Figure 1

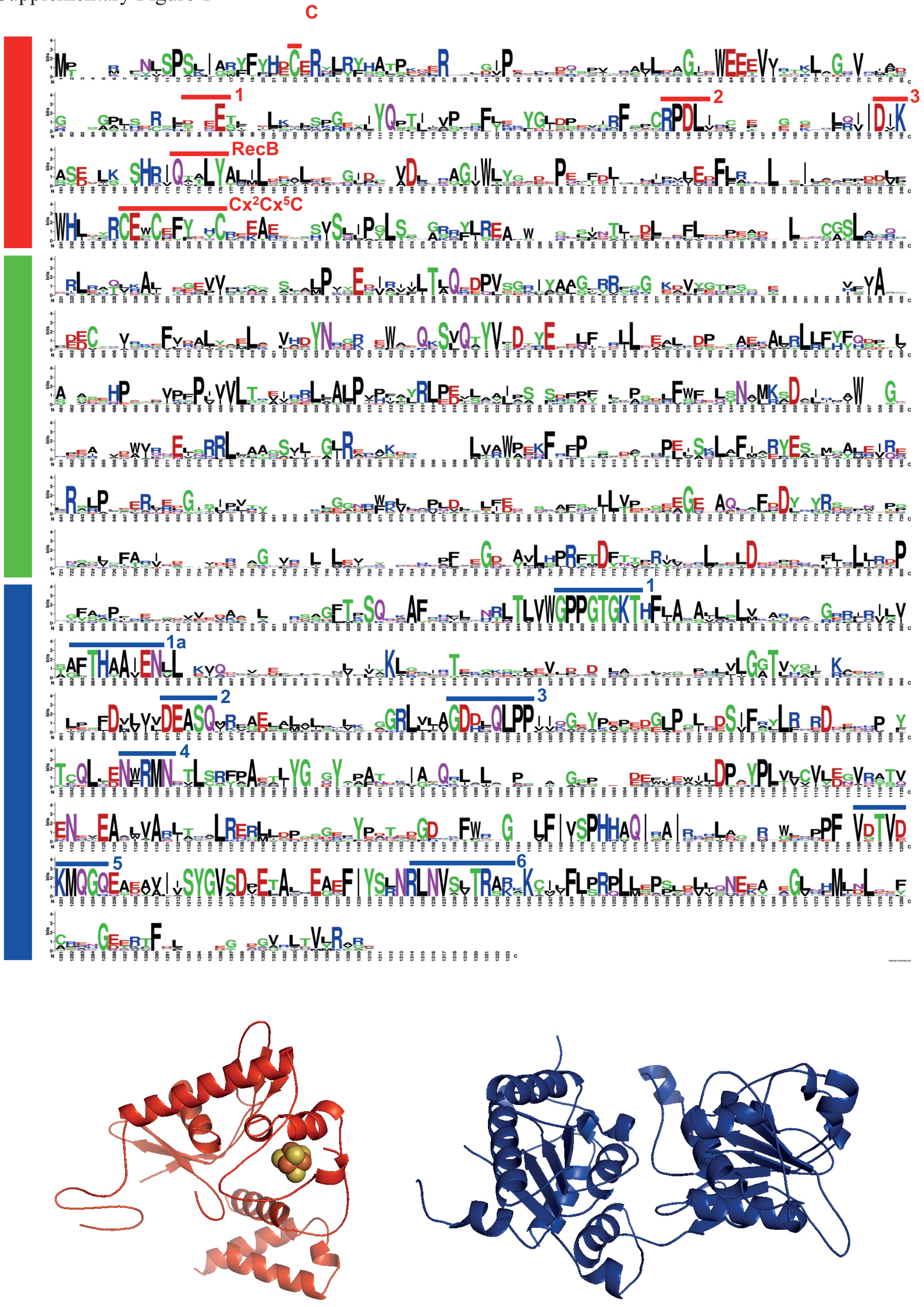

Supplementary Figure 2

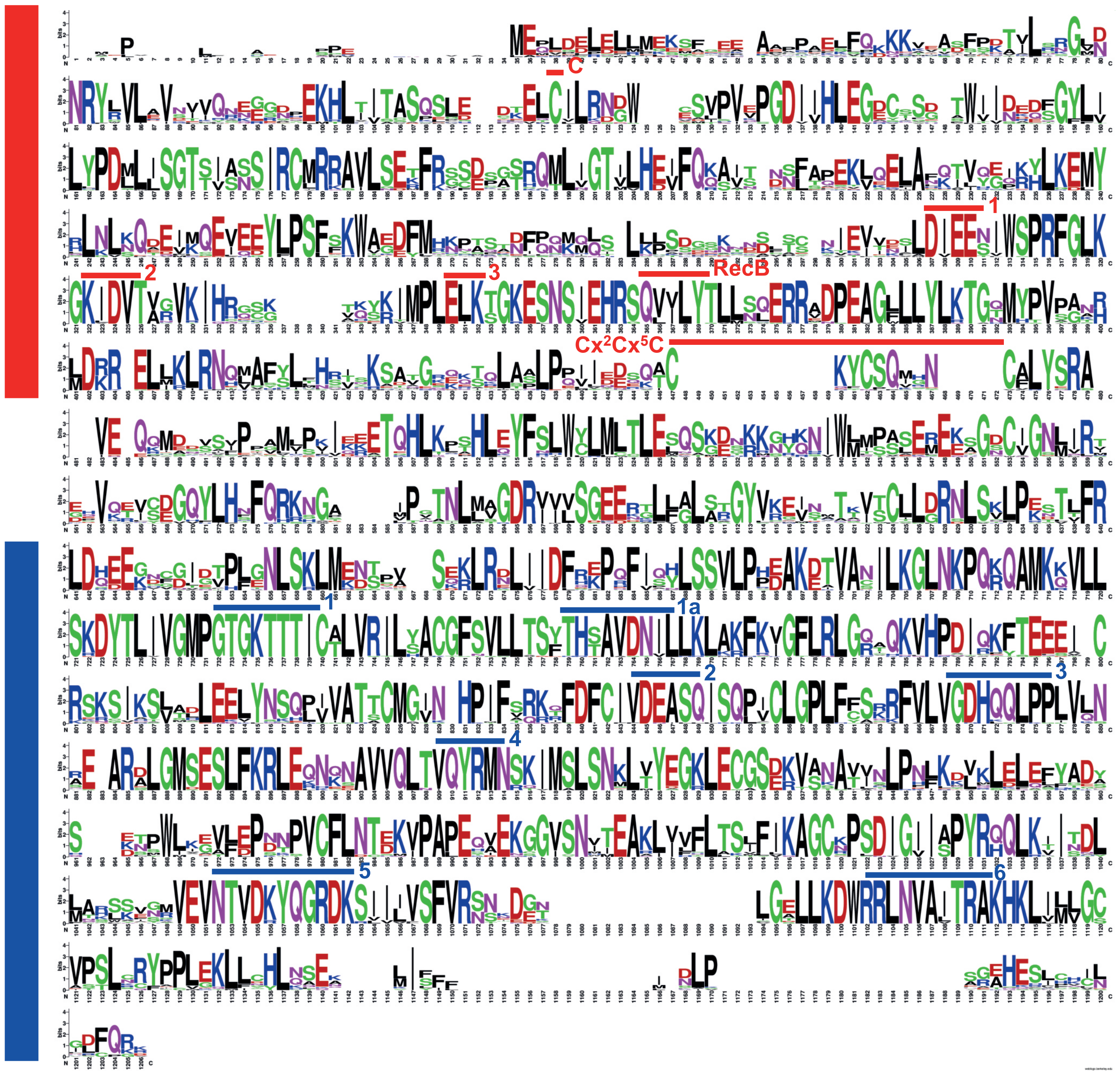

Supplementary Figure 3

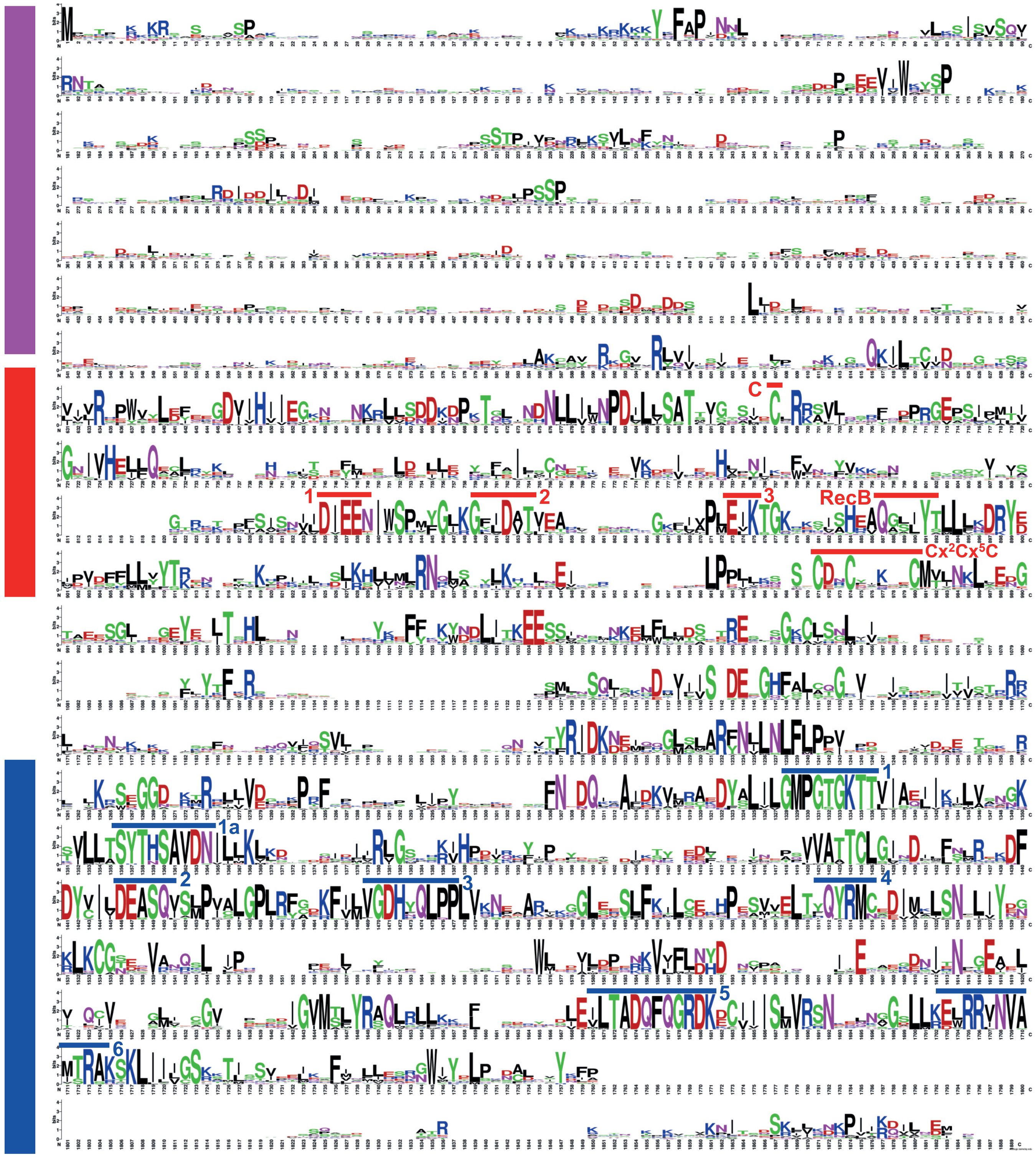

Supplementary Figure 4

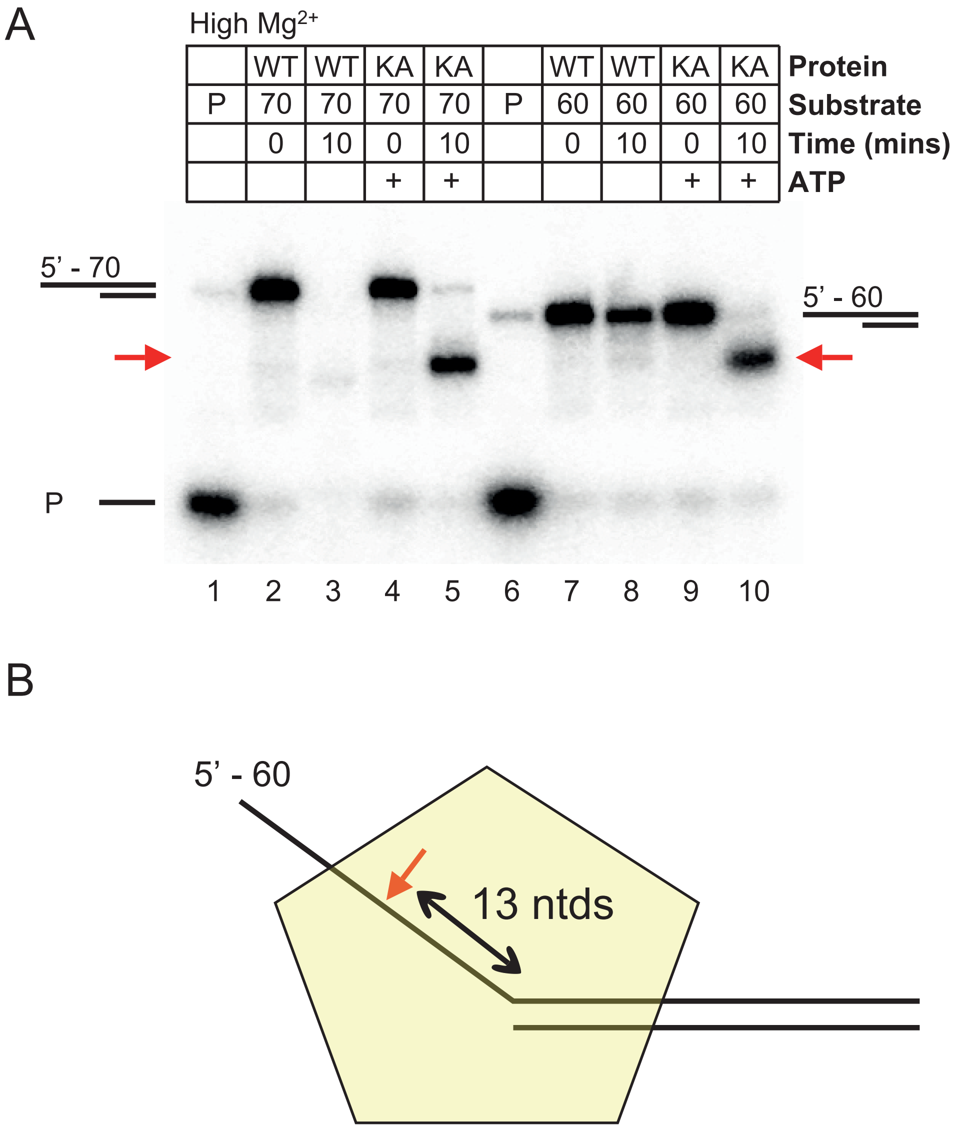

Supplementary Figure 5

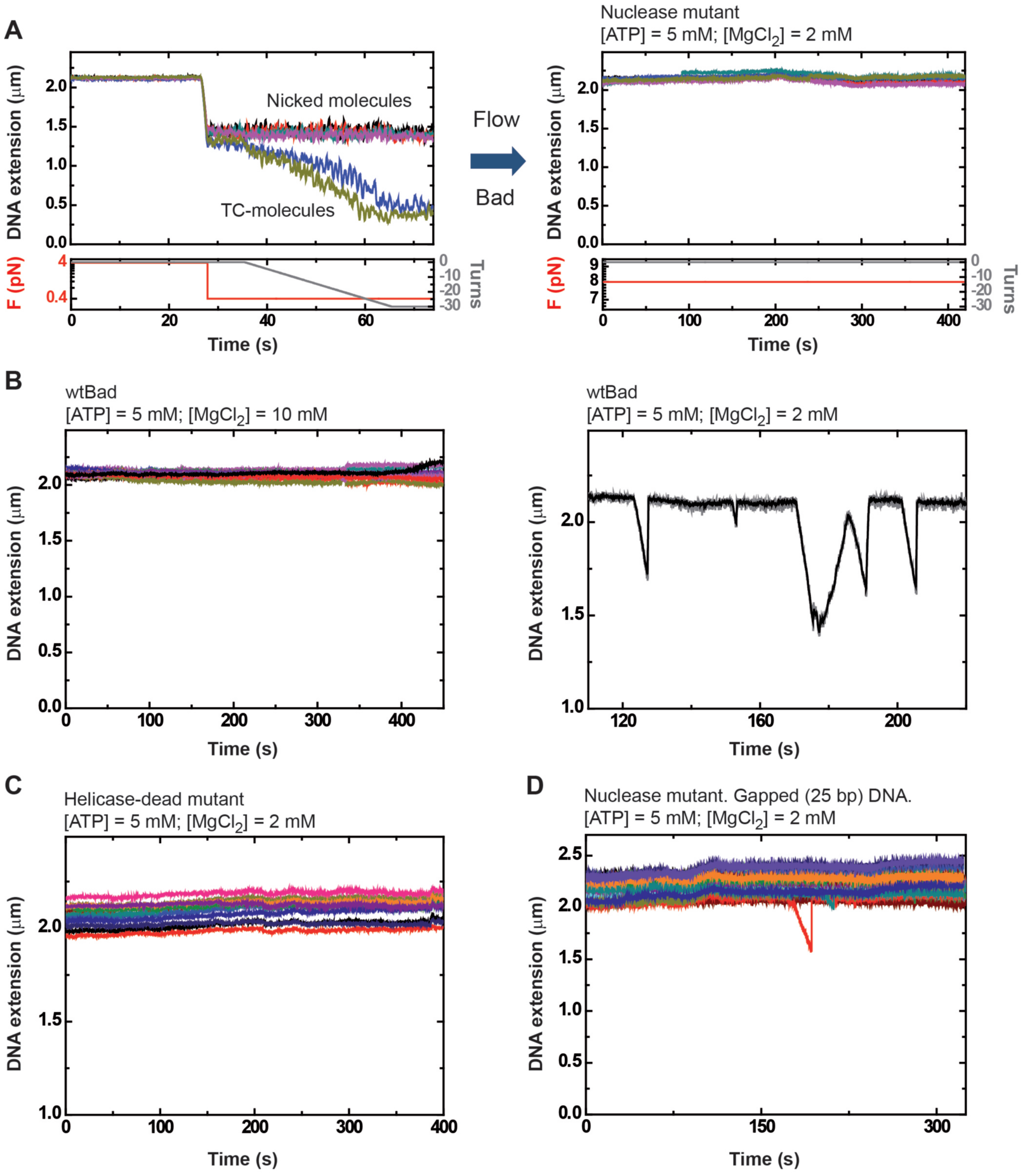

Supplementary Figure 6

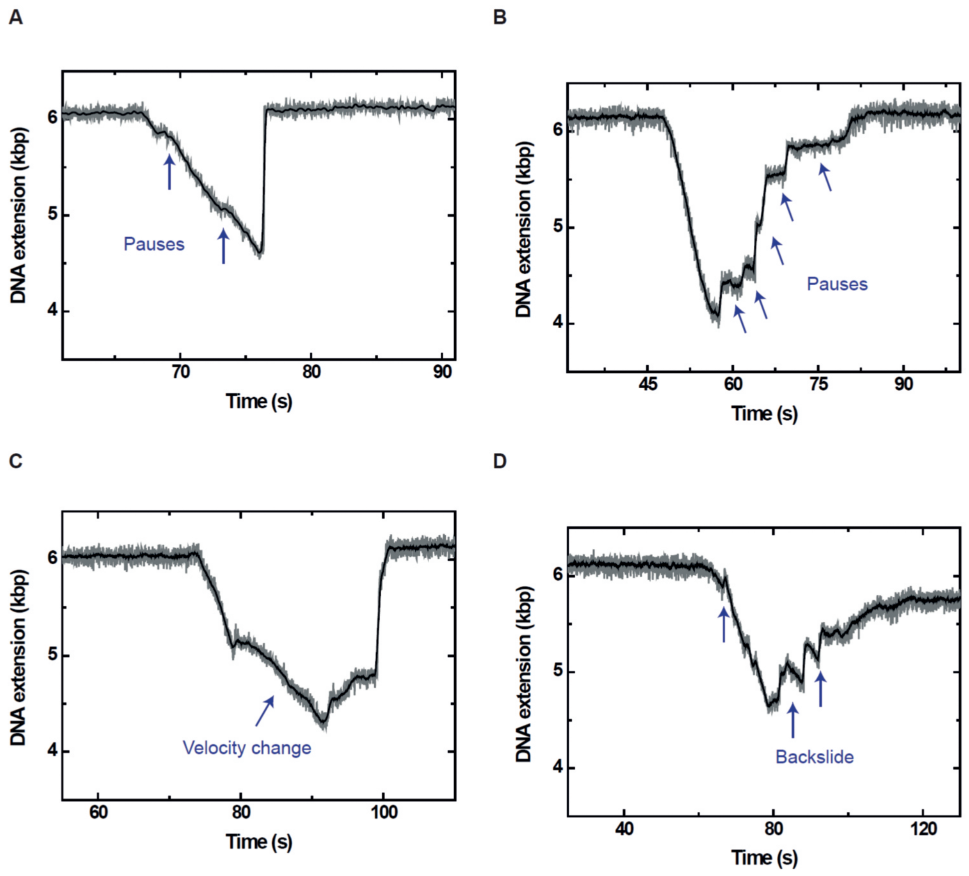

Supplementary Figure 7

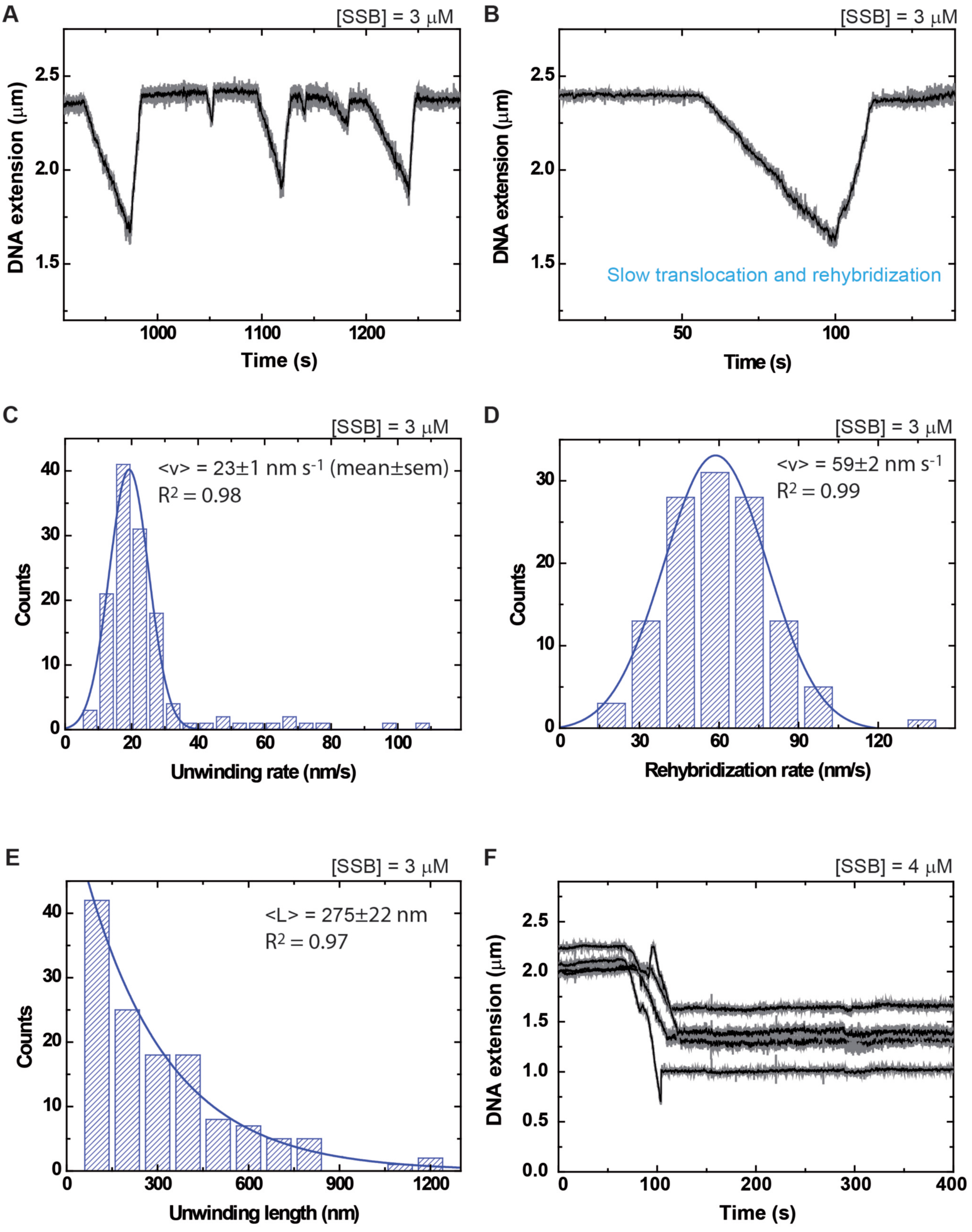
